# Cryptic diversity shapes coral symbioses, physiology, and response to thermal challenge

**DOI:** 10.1101/2024.07.09.602709

**Authors:** Hannah E. Aichelman, Brooke E. Benson, Kelly Gomez-Campo, M. Isabel Martinez-Rugerio, James E. Fifer, Laura Tsang, Annabel M. Hughes, Colleen B. Bove, Olivia C. Nieves, Alyssa M. Pereslete, Darren Stanizzi, Nicola G. Kriefall, Justin H. Baumann, John P. Rippe, Plinio Gondola, Karl D. Castillo, Sarah W. Davies

## Abstract

Coral persistence in the Anthropocene is shaped by interactions among holobiont partners (coral animals, microbial symbionts) and their environment. Cryptic coral lineages–genetically distinct yet morphologically similar groups–are critically important as they often exhibit functional diversity relevant to thermal tolerance. Additionally, environmental parameters such as thermal variability may promote tolerance, but how variability interacts with holobiont partners to shape responses to thermal challenge remains unclear. Here, we identified three cryptic lineages of *Siderastrea siderea* in Bocas del Toro, Panamá that differ in distributions across inshore and offshore reefs, microbial associations, holobiont phenomes, and skeleton morphologies. A thermal variability experiment failed to increase thermal tolerance, but subsequent thermal challenge and recovery revealed one lineage maintained elevated energetic reserves, photochemical efficiency, and growth. Lastly, coral cores highlighted that this lineage also exhibited faster growth historically. Functional variation among cryptic lineages highlights their importance in predicting coral reef responses to climate change.

**Teaser:** Cryptic host diversity drives coral phenotypes relevant to climate change.

## Introduction

Climate change is altering environments at unprecedented rates, resulting in warmer and increasingly variable environments with more extreme events (*1*). An organism’s response to such rapid changes (*e.g.,* through shifts in thermal limits, *2*) is influenced by their genetic background, environment, and interactions between these two forces (GxE; *2*, *3*). Understanding and predicting the relative importance of these factors on fitness is fundamental as environments continue to change, species ranges shift, and localized extinctions occur (*1*, *2*).

Coral reefs represent one of the most productive and economically valuable ecosystems (*4*, *5*) threatened by global (*e.g.,* warming, acidification) and local (*e.g.,* nutrient pollution, overfishing) stressors (*6*). These stressors have increased the frequency and severity of coral bleaching–loss of the coral’s obligate symbiotic algae (*7*)–which is projected to worsen under current emissions trajectories (*8*). However, reef environments are not changing homogeneously, and while our understanding of which reefs and species are more bleaching resistant is advancing (*9*), predicting their future remains challenging due to complexities governing coral resilience, including environmental variation, host genetics, and associations with diverse algal and bacterial symbionts (*10*).

Coral ‘holobionts’ encompass complex symbioses among coral hosts, algal symbionts (Symbiodiniaceae), and a diverse array of microorganisms, all interacting to shape aggregated holobiont phenotypes (*i.e.,* phenomes). Each member of the holobiont contributes to coral bleaching heterogeneity, including genetic variation of the host (*11*), algal symbiont communities (*12*), and bacterial communities (*13*). For coral hosts, genomic sequencing has revealed a surprising level of cryptic diversity, including cryptic lineages (*i.e.,* distinct genetic clusters previously characterized as one species, *14*, *15*). These lineages can also interact with a diversity of holobiont members to produce functionally distinct phenotypes. For example, a lineage in the *Acropora hyacinthus* species complex more frequently hosts *Durusdinium* algae and exhibits higher thermal tolerance, even though it coexists with other lineages on the same reef (*16*). Together, interactions among these cryptic lineages and holobiont members likely play a role in determining bleaching outcomes.

In addition to holobiont genetic variation, environmental heterogeneity can influence coral bleaching patterns (*17*) and a growing body of literature links coral thermotolerance to high frequency temperature variability, also termed diel thermal variability (DTV) (*e.g., 18*, *19*). DTV is theorized to “prime” (*i.e.,* beneficial acclimation hypothesis, *20*) organisms to more effectively respond to and recover from heat stress (*21–23*). However, DTV is correlated with other environmental parameters that can also produce beneficial acclimatory effects (*e.g.,* light, flow, *24*, *25*), and it remains unclear whether this variability facilitates thermotolerance via priming and/or whether environmental selection in high DTV environments is selecting for more thermally tolerant individuals. For example, cryptic coral lineages are known to exhibit divergent spatial distributions across depths (*26*), which likely involves adaptations in photosynthetic pigment concentrations and skeletal traits that can have important effects on light harvesting potential (*27*, *28*). Additionally, lineages and their algal symbionts (*29*) have been shown to segregate across inshore-offshore gradients, where offshore habitats experience lower turbidity, higher flow, and more stable temperatures relative to inshore habitats (*e.g., 30*). Together, these patterns of complex environmental heterogeneity likely produce adaptive phenotypes through a combination of acclimation and selection for unique holobiont combinations.

To investigate the influence of coral holobiont diversity and environmental history on coral phenomes, we characterized holobiont genetic diversity of the reef-building coral *Siderastrea siderea* from three inshore and three offshore sites in the Bocas del Toro reef complex (BTRC), Panamá. We discovered three cryptic lineages with unique symbiotic associations that differed in their distributions across the seascape, as well as distinct baseline phenomes and skeletal morphology. Next, we conducted a 50-day DTV experiment, followed by thermal challenge and recovery to test the hypothesis that exposure to DTV would increase coral resistance to thermal challenge. In contrast to our hypothesis, we found that cryptic lineages differed in their response to thermal challenge, while the effect of experimental DTV treatment was minimal beyond promoting growth. Interestingly, lineages also differed in their growth, especially under thermal challenge. To determine whether these growth differences between lineages were conserved *in situ*, we used coral cores to contrast long term growth trajectories between lineages. We found that these records were consistent with experimental patterns, with lineages differing in skeletal density and linear extension rates. Together, these data showcase the strong influence that cryptic lineages have in shaping coral distributions, symbioses, thermotolerance, and growth. Failure to appreciate this genetic diversity will ultimately lead to challenges when predicting coral responses to climate change.

## Results

### Three lineages of *Siderastrea siderea* with distinct distributions and symbioses found in Bocas del Toro

This study focused on the ubiquitous Caribbean reef-building coral, *Siderastrea siderea,* collected primarily from six sites in the BTRC, Panamá across inshore (Punta Donato=PD, STRI Point=SP, Cristobal Island=CI) and offshore (Bastimentos North=BN, Bastimentos South=BS, Cayo de Agua=CA) sites (Fig. 1A, N=9 corals/site, Table S1A). Corals from an additional inshore (Punta Laurel=PL) and offshore (Drago Mar=DM) site were included for historical growth data (Fig. 1A), which is described below. 2b-RADseq genotyping was conducted on 54 coral colonies. Admixture ancestry of individuals across sites (Fig. 1A; Fig. S1A), principal coordinate analysis (PCoA) based on the identity by state (IBS) matrix (Fig. S1B), and hierarchical clustering of pairwise IBS values (Fig. S1C,D), supported the presence of three distinct coral host genetic clusters (hereafter referred to as cryptic lineages L1, L2, and L3) across BTRC. Pairwise global weighted F_ST_ values revealed high divergence between these genetic clusters (L1 vs L2=0.17, L1 vs L3=0.18, L2 vs L3=0.12; Fig. S1B), suggesting they represent three cryptic lineages of a *S. siderea* species complex. Cryptic lineages differed in their spatial distributions across BTRC, with more L1 individuals sampled at offshore sites (83%; 24/29 offshore/total L1 individuals) and more L2 individuals sampled at inshore sites (94%; 17/18 inshore/total L2 individuals; *X*^2^=23.57, *p*<0.001). L3 was the least abundant lineage, with only three individuals observed at SP (Fig. 1A). CI is the only site where two lineages were sampled in equal proportion (n=4 L1, n=4 L2; Fig. 1A). While ADMIXTURE results at K=2 suggest that L3 individuals are of mixed ancestry between L1 and L2, L3 fully resolves as a distinct lineage at K=3 (Fig. S1A). Cryptic lineages had low admixture, and the individual with the most admixture had <5% assigned to a second ancestral population. Given the small sample size for L3, these individuals were excluded from all downstream analyses.

**Figure 1.**
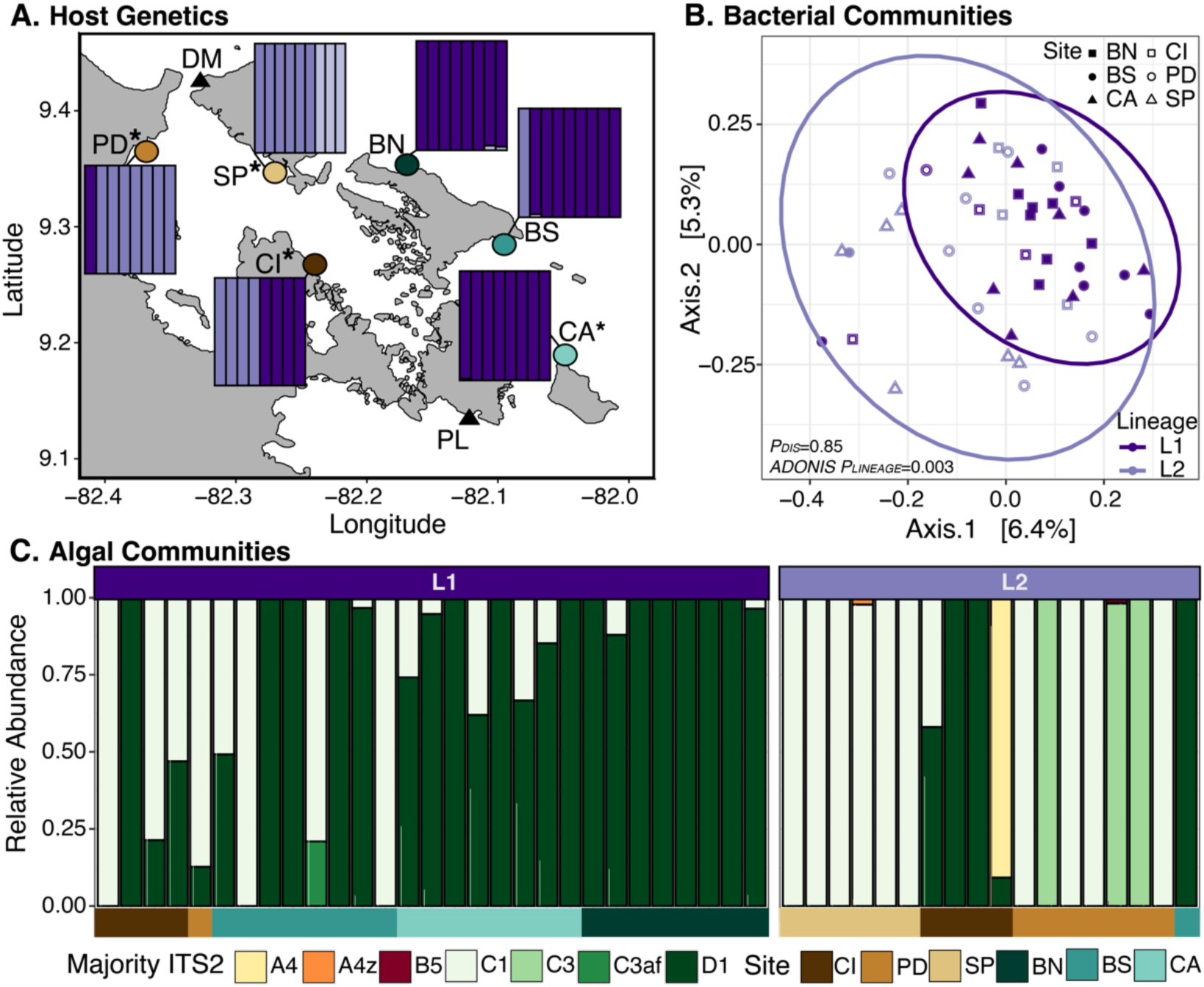
Cryptic lineages exhibit distinct symbioses. **A.** Map indicating eight sites across the Bocas del Toro reef complex in Panamá where *Siderastrea siderea* colonies were collected. Primary sites include three inshore (brown shades: PD=Punta Donato, SP=STRI Point, CI=Cristobal Island) and three offshore (green shades: BN=Bastimentos North, BS=Bastimentos South, CA=Cayo de Agua) sites. Coral cores were taken from these six sites plus an additional two sites (black triangles; inshore=Punta Laurel=PL, offshore=Drago Mar=DM). Asterisks indicate sites where temperature loggers were recovered. ADMIXTURE results from 2b-RADseq data for K=3 are grouped by collection site. Columns represent an individual and the bar color shows assignment to one of three ancestral populations (L1: dark purple, L2: medium purple, L3: light purple). **B.** Bray-Curtis dissimilarity principal coordinate analysis (PCoA) of coral bacterial communities using all ASVs from cleaned and rarefied 16S metabarcoding data. Ellipses represent 95% confidence intervals, *ADONIS P*-values indicate significant community differences, and *P_DIS_* values compare dispersion across lineages. **C.** Bar plots of Symbiodiniaceae majority ITS2 sequence relative abundance data, colored by majority ITS2 type. Each column of the bar plot represents a host colony, with color blocks representing site of origin, as in **A**.

16S metabarcoding of the same coral colonies determined that bacterial communities were significantly different between L1 and L2 (Fig. 1B; *ADONIS p=*0.003; Table S2) when sampled before experimental acclimation (hereafter termed ‘baseline’, full experimental design detailed in Fig. S2). No differences were observed for any bacterial community alpha diversity metrics (ASV richness, Shannon’s index, Simpson’s index, and evenness) across host lineages for samples at baseline (Fig. S3; Table S3).

ITS2 metabarcoding revealed that when Symbiodiniaceae communities were aggregated by the majority ITS2 sequence assigned by SymPortal (*31*), 19 Symbiodiniaceae ITS2 defining intragenomic variants (DIVs) matched the C1 majority ITS2 sequence (*Cladocopium goreaui*), two DIVs matched C3, nine DIVs matched D1 (*Durusdinium trenchii*), and one DIV matched each of the A4, A4z, B19, B5, and C3af majority ITS2 sequences. At baseline, significantly more L1 (72.4%) hosted >50% relative abundance of *D. trenchii* relative to L2 (22.2%) (Kruskal-Wallis *X*^2^=12.68, *p*=0.0004). When only considering corals from the site where an equal proportion of L1 and L2 individuals were collected (CI) at baseline, differences in the relative abundance of *D. trenchii* between lineages were no longer significant, with 25% of L1 and 75% of L2 hosting >50% relative abundance of *D. trenchii* (Kruskal-Wallis *X*^2^=0.788, *p*=0.37).

### Cryptic host lineages exhibited distinct phenomes and skeleton morphologies

Corals of L1 and L2 ancestry exhibited distinct holobiont phenomes at baseline (Fig. 2A; *ADONIS p*<0.0001, partial Omega-squared (𝜔^2^)=0.14; Table S4). Lineages were distinguished by the first principal component (PC), with loadings for energy reserves (symbiont density, host and symbiont carbohydrate, chlorophyll *a*, and protein) positively correlated with L1 (Fig. 2A). Comparing holobiont phenomes (Fig. 2A) with individual physiology results (Fig. S4, Table S5) supports L1 having significantly greater total protein, total host and symbiont carbohydrates, chlorophyll *a*, and symbiont density relative to L2 at baseline.

**Figure 2.**
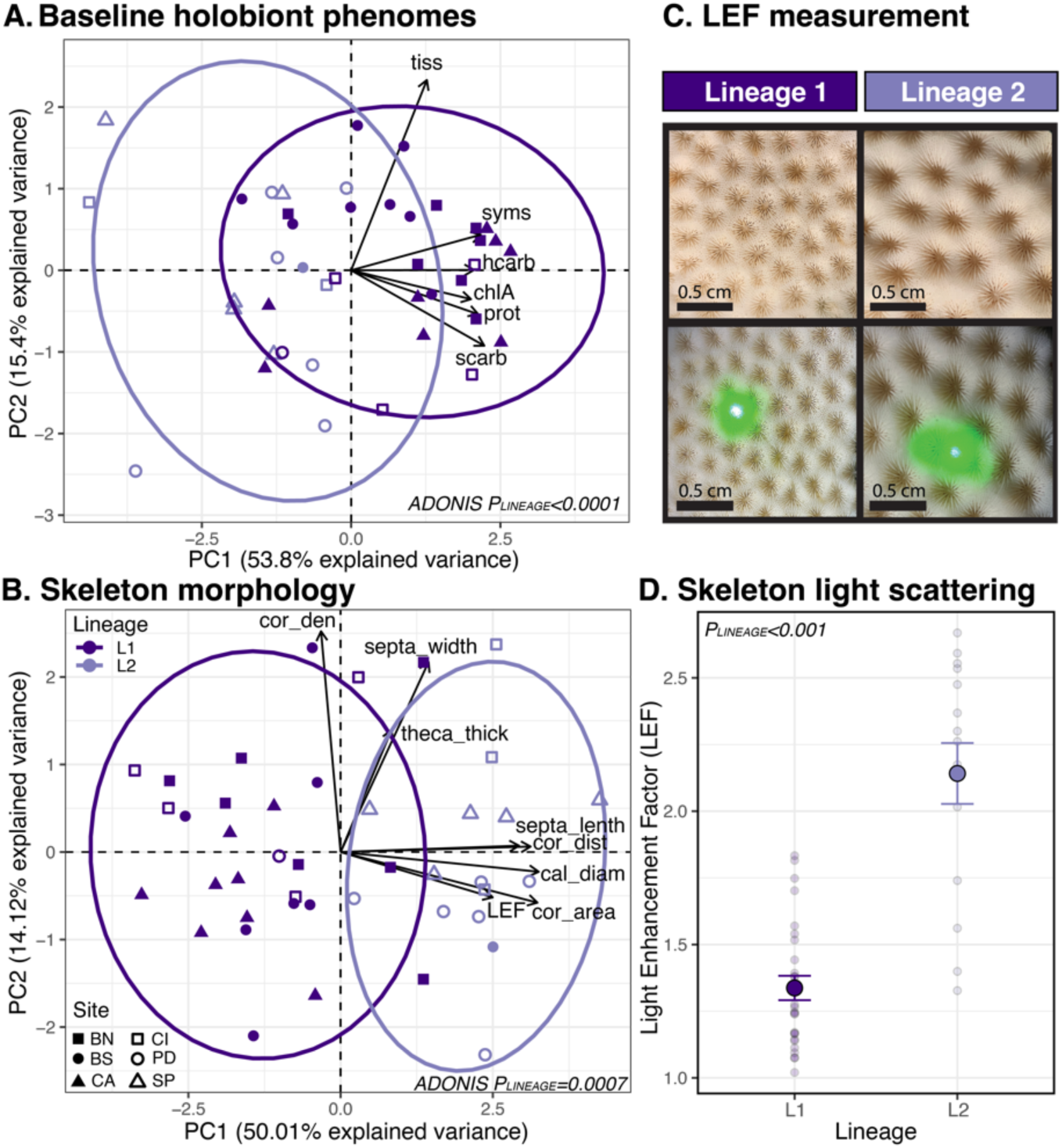
Phenomes and skeleton morphology differ between cryptic host lineages. **A.** Principal component analysis (PCA) of *log*-transformed holobiont phenomes showcasing differences between L1 and L2 at baseline. Phenotypes include tissue thickness (tiss; mm), symbiont density (syms; cells cm^-2^), host and symbiont carbohydrate (hcarb and scarb, respectively; mg cm^-2^), chlorophyll *a* (chlA; ug cm^-2^), and total protein (prot; mg cm^-2^). P-value indicates significant difference in holobiont phenomes determined from a PERMANOVA, only individuals with data for all phenotypes were included (N=42; Table S4). **B.** PCA of *log*-transformed coral skeleton morphology sampled at baseline. Phenotypes include corallite area (cor_area; cm^-2^) corallite density (cor_den; # cm^-2^), septal width (septa_width; cm), theca thickness (theca_thick; cm), distance between corallites (cor_dist; cm), septal length (septa_len; cm), calyx diameter (cal_diam; cm), and light enhancement factor (LEF). Statistical analyses (Table S6) and sample size as in **A**. **C.** Top: photos of representative skeletons from L1 (left) and L2 (right). Bottom: photos of light enhancement factor (LEF) measurements, illustrating lineage differences in the ability for the skeleton to enhance the light field for algal photosynthesis. **D.** Lineage differences in skeleton LEF (L1: N=27, L2: N=15). P-value indicates significant difference in LEF (Table S7). Large points represent mean LEF ± standard error for each lineage, smaller points represent an individual fragment’s LEF.

Corals of L1 and L2 ancestry exhibited distinct skeleton morphologies at baseline (Fig. 2B). Principal Component Analysis (PCA) of all skeletal parameters showcased that combined skeleton morphologies were significantly different across lineages (Fig. 2B; *ADONIS p*=0.0007, 𝜔^2^=0.18; Table S6). Lineages were distinguished by the first PC, with loadings for septa length, distance between corallites, calyx diameter, corallite area, and light enhancement factor (LEF) positively correlated with L2 (Fig. 2B). L2 had larger corallite area when compared to L1 (58% larger; *p*<0.0001; Fig. S5A; Table S7). This pattern was consistent even at CI where L1 and L2 co-occur, with L2 (N=4) maintaining larger corallites than L1 (80% larger; N=4; Fig. S5B; *p*=0.001; Table S7). This variation in skeletal morphology contributed to a significantly higher LEF in L2 compared to L1 (Fig. 2C,D; *p*<0.0001; Table S7), indicating an increased ability of L2 skeletons to enhance the light field for algal photosynthesis. Similarly, L2 maintained significantly greater LEF at CI, where lineages co-occur (Fig. S5C; *p*=0.04; Table S7).

### Diel thermal variability influenced growth and bacterial communities, but host lineages shaped algal symbioses

To explore how lineages collected from sites across BTRC respond to differences in diel thermal variability (DTV), we conducted a 50-day DTV experiment where corals were exposed to either no variability control conditions (average daily mean ± average daily variability=29.5 ± 0.4°C) or DTV (29.4 ± 2.9°C; treatment temperatures summarized in Table S1B). Buoyant weight assessments showed an effect of DTV on growth, with corals exposed to DTV growing faster than control corals (32.9% increase; Fig. 3A; *p*=0.02; Table S8). Additionally, under DTV, L1 growth trended towards being faster than L2 (36.9% increase; Fig. 3A; *p*=0.08).

**Figure 3.**
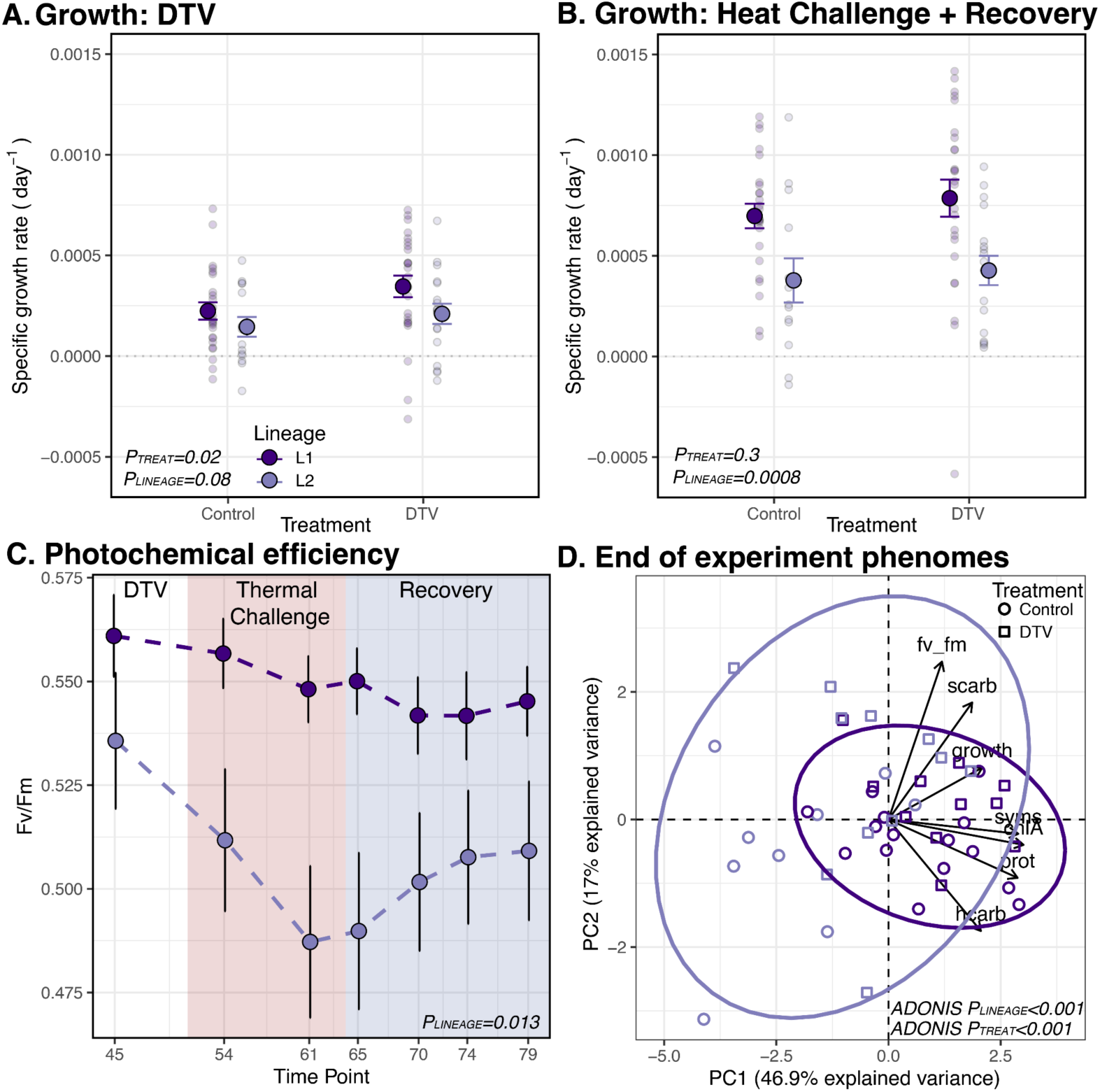
Lineage responses to diel thermal variability (DTV) and thermal challenge. Specific growth rate day^-1^ between DTV treatments showing lineage growth throughout the DTV experiment (**A**) and thermal challenge and recovery (**B**). Large points represent mean growth ± standard error for each lineage, and smaller points represent an individual fragment’s growth. P-values indicate significant differences in growth (Table S8). **C.** Photochemical efficiency of photosystem II (Fv/Fm) across seven time points including the end of DTV treatment, and throughout heat stress (red shaded) and recovery (blue shaded). Points represent mean Fv/Fm ± standard error for each lineage (Table S9). **D.** Principal component analysis (PCA) of *log*-transformed holobiont phenomes at the end of the experiment. Phenotypes include specific growth rate through 50 days in DTV (growth), total protein (prot; mg cm^-2^), host and symbiont carbohydrate (hcarb and scarb, respectively; mg cm^-2^), chlorophyll *a* (chlA; mg cm^-2^), symbiont density (syms; cells cm^-2^), and Fv/Fm. Only individuals with data for all phenotypes were included (N=46), and P-values indicate significant differences in holobiont phenomes determined from PERMANOVA (Table S4).

At the end of the DTV experiment, coral fragments were subsampled for 16S and ITS2 metabarcoding to explore how DTV modulated symbioses. 16S data showed that bacterial communities were no longer structured by host lineage (*ADONIS p*=0.11; Fig. S6A). However, communities in DTV were distinct from those in control (*ADONIS p*=0.045; Fig. S6B). Additionally, ASV richness was higher in control corals compared to corals exposed to DTV (Fig. S6I; *p*=0.023; Table S3). No other differences were observed for any bacterial community diversity metrics (ASV richness, Shannon’s index, Simpson’s index, and evenness) across host lineage or variability treatment (Fig. S6 C-J; Table S3). ITS2 data showed that Symbiodiniaceae communities continued to be differentiated by cryptic host lineage after 50 days of DTV (Fig. S7) with 49.1% of L1 hosted >50% relative abundance of *D. trenchii* compared to 12.9% in L2 (Kruskal-Wallis *X*^2^=16.82, *p*<0.0001).

### Host lineage, not DTV, drove responses to thermal challenge, but both shaped holobiont phenomes

To explore how DTV influenced response to thermal challenge and recovery, after 50 days of DTV all corals were exposed to a two-week thermal challenge followed by a two-week recovery. Buoyant weights showed that growth benefits afforded from exposure to variability were not sustained and prior exposure to DTV failed to rescue growth during the thermal challenge and recovery periods (Fig. 3B; *p*=0.3; Table S8). However, L1 grew significantly faster than L2 during these periods (45.4% increase; Fig. 3B; *p*=0.0008; Table S8) and L1 maintained higher photochemical efficiency of photosystem II (Fv/Fm) than L2 throughout thermal challenge and recovery (7.8% increase; Fig. 3C; *p*=0.013; Table S9). Prior exposure to DTV did not have a significant effect on Fv/Fm during thermal challenge and recovery (Fig. 3C; *p*=0.21; Table S9). Together, these data suggest that lineages exhibit divergent responses to thermal challenge.

At the end of the thermal challenge and recovery periods, coral fragments were flash frozen and holobiont phenomes were examined. Similar to baseline conditions (Fig. 2A), host lineages maintained distinct holobiont phenomes at the end of the experiment (Fig. 3D; *ADONIS p*<0.001; 𝜔^2^=0.201; Table S4), with loadings for host and symbiont energy reserves, symbiont density, and chlorophyll *a* positively correlated with L1. Individual physiology metrics (Fig. S4, Table S5) demonstrate that L1 maintained significantly more total protein, total host carbohydrates, chlorophyll *a*, and symbiont density than L2 at the end of the experiment. Additionally, DTV treatment influenced holobiont phenomes, with control corals distinguishing from corals exposed to variability (Fig. 3D; *ADONIS p*<0.001; 𝜔^2^=0.177; Table S4). Individual physiology metrics suggest that this difference was driven by greater symbiont density and symbiont carbohydrates, but reduced host carbohydrates, in corals exposed to DTV relative to control (Fig. S4).

### Host lineages exhibit distinct patterns of historical growth

Lastly, to explore whether the lineage differences in growth observed under experimental conditions are conserved *in situ*, we assessed patterns of annual growth from coral cores between 1980-2014 that were genotyped via 2b-RADseq as described above (L1: N=16, L2: N=8 cores). L1 maintained significantly higher linear extension (*p*=0.03; Fig. 4A) and lower skeletal density (*p*=0.004; Fig. 4B) compared to L2 (Table S10). While L1 cores trended towards greater calcification than L2, these differences were not statistically significant (*p*=0.11; Fig. 4C; Table S10). These patterns were consistent when considering linear extension, skeletal density, and calcification data across the entire history of the coral cores, with the longest record being 1880-2014 (Fig. S8; Table S10). While both annual mean and annual summer mean temperatures have increased in the BTRC over the lifetime of the coral cores analyzed (Fig. S9A,B; *p<*0.001; Table S11), calcification was not influenced by temperature across any time interval (Fig. S9C-F; Table S12). However, L1 did maintain significantly greater calcification across both annual mean and annual summer mean temperatures for both time intervals considered (all *p*<0.001; Fig. S9C-F; Table S12).

**Figure 4.**
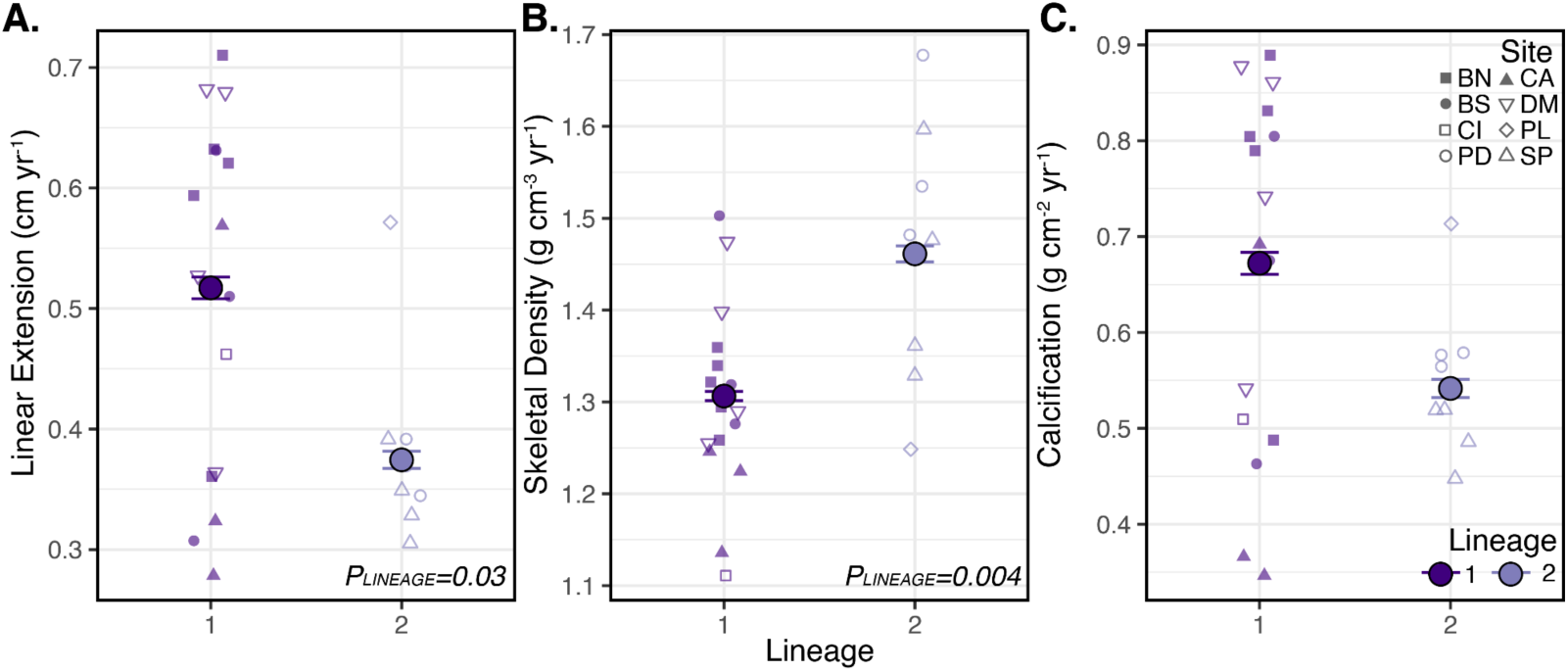
Cryptic host lineages differ in historical growth. Linear extension (**A**), skeletal density (**B**), and calcification (**C**) across cryptic host lineages. For all panels, large points represent the mean ± standard error of each metric and smaller points represent an individual coral core. Shape of the smaller points represents the site of origin for that core (Offshore: BN=Bastimentos North, BS=Bastimentos South, CA=Cayo de Agua, DM=Drago Mar; Inshore: PD=Punta Donato, SP=STRI Point, CI=Cristobal Island, PL=Punta Laurel). Only data in the interval of 1980-2014 are included here (N=16 L1 cores, N=8 L2 cores; Table S10).

## Discussion

Understanding how holobiont diversity is partitioned across the seascape is critical to predicting coral responses to climate change (*11*, *12*). Here, we identified three cryptic host lineages (L1, L2, L3) in a *Siderastrea siderea* species complex across BTRC that varied in baseline phenotypes including algal and bacterial symbiont communities, energetic reserves, and skeletal morphology relevant to their *in situ* light environment. Additionally, L1 corals maintained elevated energetic reserves throughout the experiment, as well as elevated photochemical efficiency and growth throughout subsequent thermal challenge and recovery. Finally, coral core data sampled from lineages across BTRC demonstrated that these distinct growth patterns observed between lineages in the experiment were consistent with patterns of historical growth.

This work builds on the growing evidence for widespread cryptic diversity in corals (*15*), which has been detected at large (*e.g.,* archipelago-wide, *32*) and small spatial scales, including within reefs in Puerto Rico (*26*), the Florida Keys (*33*), and American Samoa (*16*). Depth has emerged as a common driver of lineage differentiation in corals, with relevant abiotic factors including temperature, seawater optical properties (*26*, *33*), and small-scale current patterns (*34*). These cryptic lineages have also been shown to exhibit divergence in ecologically relevant traits. For example, sympatric lineages of the *Pachyseris speciosa* species complex differ in skeletal morphology and holobiont physiology (*27*). Across larger spatial scales, only one of three *Acropora hyacinthus* cryptic lineages was able to occupy habitats along a range expansion front in Japan, which was attributed to divergence at loci associated with adaptation to temperate, seasonally fluctuating environments (*32*). Thermal variability has also been associated with differential distributions of *A. hyacinthus* cryptic lineages that exhibit distinct thermal tolerances in American Samoa (*16*, *35*). Here, we find that cryptic lineages vary in ecologically relevant phenomes (*i.e.,* energetic reserves, growth) and in their response to thermal challenge.

We find evidence for environmental structuring of lineages and that unique skeleton morphologies could contribute to their success in these distinct environments. The cryptic host lineages identified here were structured across an inshore to offshore gradient in BTRC, with L2 and L3 more prevalent inside Bahia Almirante (inshore), and L1 more prevalent outside the bay (offshore). Inshore BTRC sites are characterized by limited influence from the open ocean, riverine inputs that deliver nutrients, agricultural run-off and sewage to the bay, higher turbidity, and most recently hypoxic events that have altered coral communities (*36*, *37*). We find evidence that lineages exhibit unique skeleton morphologies that could contribute to their success in these distinct environments. Namely, L2 skeleton morphology suggests that this lineage is better adapted to the low light environments of inshore BTRC to promote algal photosynthesis. The ability of L2 skeletons to better amplify incoming light could also explain its lower symbiont densities and chlorophyll *a* concentrations compared to L1, which would serve as a mechanism to reduce pigment packing while “sensing more light” (*28*, *38*). These skeleton morphology differences persisted even when lineages co-existed in the same environment (CI), which suggests a genetic basis for this trait. Future explorations of this system would benefit from collecting precise colony depths in addition to diffuse attenuation coefficients for downwelling irradiance (K_d_) from each site to link *in situ* light environments with cryptic lineage assignment. This information, along with additional optical traits of the coral holobiont, particularly metrics taken with intact tissue, would provide more detailed information on how seawater optical properties drive lineage distributions in BTRC (*e.g.*, *39*).

These cryptic lineages not only occupied distinct habitats, they also exhibited unique symbioses. L1 and L2 hosted distinct communities of bacterial symbionts at baseline; however, these differences did not persist after a 50-day DTV treatment. Instead, bacterial communities of corals exposed to variability were distinct from corals in control conditions. While previous work has highlighted that bacterial communities can contribute to coral thermal tolerance (*13*), this does not seem to be the main driver here given that we observed thermal tolerance differences among corals with similar bacterial communities. In contrast, algal communities were strongly structured by lineage, with L1 associating with higher proportions of *Durusdinium trenchii* and hosting a unique *Cladocopium* (C3) DIV. Increased *D. trenchii* in L1 could contribute to their elevated Fv/Fm throughout thermal challenge relative to L2, as *D. trenchii* has been shown to confer thermal tolerance to their hosts (*e.g.*, *12*, *40*). Interestingly, Rose et al. (*16*) also demonstrated that a more bleaching-resistant *Acropora hyacinthus* cryptic lineage hosted greater proportions of *D. trenchii*. Here, while some L2 colonies were dominated by *D. trenchii,* this only occurred in low frequencies at sites where both L1 and L2 were sampled (CI and BS). Regardless, we posit that the elevated light enhancement factor (LEF) of L2 skeletons coupled with their acclimation to lower light environments of inshore BTRC and lower symbiont densities could have led to reduced Fv/Fm during thermal challenge. Future work investigating the impact of acclimation to different light levels on heat tolerance in these lineages is warranted. Additionally, as coral bacterial communities are strongly influenced by their host traits and environment (*41*), more thorough sampling across sites, cryptic lineages, and regions within a coral colony (*e.g., 42*) will be necessary to better identify any lineage-specific bacterial taxa.

We initially hypothesized that DTV would shape coral phenomes to increase thermal resilience (*e.g., 18*, *21*, *23*, *43*). While an effect of experimental DTV on holobiont phenomes was observed, it was largely driven by differences in growth across treatments. DTV increased growth, suggesting that it represents a promising coral restoration tool to improve growth in nursery settings (as in *43*). However, experimental exposure to DTV did not facilitate the maintenance of Fv/Fm during subsequent thermal challenge. It is possible that the DTV treatment used here was insufficient to “prime” corals (reviewed in *44*). Indeed, unique DTV temperature regimes and timing of exposure have previously resulted in variable phenotypic outcomes for the corals *Montipora capitata* (*21*) and *Acropora aspera* (*45*). In *M. capitata,* only two of four variability pre-exposure profiles altered gene expression and resulted in improved thermal tolerance in stress tests four months later (*21*). Additionally, in *A. aspera*, exposure to DTV in the month preceding a heat challenge had a larger, and ultimately deleterious, effect on heat tolerance compared to corals that experienced 1.5 years of preconditioning to variable temperatures (*45*). Previous work has also demonstrated that DTV can have negative effects when additional heat stress is present (*e.g., 46*), and BTRC reached ∼7 degree heating weeks in the year prior to coral collection (2015; NOAA Coral Reef Watch v3.1), which could have influenced responses to DTV.

Elevated growth of L1 compared to L2 was evident not only during the *ex situ* DTV experiment, but also in coral cores from across BTRC belonging to the same cryptic lineages. Century-scale data from cores of long-lived corals such as *S. siderea* allow investigation into the impact of long-term environmental conditions on coral growth and ultimately health. Here, cores sampled across inshore and offshore environments of BTRC show that L1 maintained greater linear extension, lower skeletal density, and trended towards greater overall calcification compared to L2. Previous coring studies on *S. siderea* demonstrate that reef environments play an important role in shaping long-term growth trajectories. Specifically, in *S. siderea* sampled from the southern Mesoamerican Barrier Reef System, forereef corals exhibited long-term declines in skeletal extension rates while nearshore and backreef coral extension rates were stable (*47*, *48*). The authors propose that resilience of nearshore and backreef corals is linked to their exposure to greater diurnal and seasonal thermal variability (*48*). In contrast, more widespread sampling of *S. siderea* cores across the entire Mesoamerican Barrier Reef System revealed declines in skeletal extension rates only for nearshore corals, which was attributed to exposure to land-based anthropogenic stressors and ocean warming (*49*). While it is possible that the lineage differences in historical growth found here are driven by reef environments, because lineage and environment were almost fully confounded in this coring sampling design, it is also possible that previous work on coral growth trajectories are complicated by the presence of *S. siderea* cryptic diversity across the Mesoamerican Barrier Reef System. Future work sampling cores from coexisting lineages would disentangle the role of environmental and genetic factors in determining long-term growth trajectories. A more thorough characterization of general environmental conditions at these sites (*e.g.,* nutrient concentrations, pH, and dissolved oxygen) is also needed, as sites where L1 and L2 are sympatric suggest other environmental characteristics could be driving differentiation and distributions of *S. siderea* cryptic lineages.

*Siderastrea siderea* is a horizontally transmitting, gonochoric broadcast spawning coral, with colonies of separate sexes spawning gametes to produce aposymbiotic larvae that spend time in the water column before settling, leading to the potential for broad population connectivity across great distances (up to 1200 km, *50*). While much more work is warranted, we propose that in BTRC, the distinct light environments across inshore and offshore reefs along with physical characteristics of the archipelago (*37*, *51*) result in spatially varying selection on cryptic lineages that are uniquely adapted to distinct light environments. While few sites were found to host multiple lineages and no site hosted all three, sampling for this study was limited to <8 m and exact depths of corals were not recorded. Therefore, we hypothesize that additional sampling across depth within individual sites will reveal differential depth distributions of the lineages that reflect patterns observed across inshore and offshore environments (*i.e.,* L1 associated with higher light and L2 associated with lower light). As was previously demonstrated by Quigley et al. (*52*), it is also likely that environmental pools of algae are much more diverse than communities hosted by adult corals, and therefore once recruits begin establishing symbiosis, algal symbionts likely compete through a “winnowing” process with dominance depending on local environmental conditions (*i.e.,* light, depth) that are further shaped by coral colony and skeleton morphology (*28*). Surviving recruits of distinct lineages then develop associations with specific Symbiodiniaceae in environments that differ in temperature, light, and nutrients, likely resulting in further acclimation to local conditions. Together, these genetic and environmental factors interact to determine the patterns of responses observed here, where unique holobiont partners shape variation in phenomes, response to thermal challenge, and historical growth. Ultimately, reciprocal transplant experiments are needed to disentangle the relative roles of adaptation and acclimation in the observed phenotypes between lineages. This work highlights the importance of understanding cryptic coral diversity when determining species responses to future climate change and in restoration planning.

## Materials and Methods

### Identification of cryptic lineages

In August 2016, nine visually healthy colonies (20-30 cm diameter) of the reef-building coral species *Siderastrea siderea* were collected between 2.5 and 8 m depth at each of six sites (54 colonies total, permit SE/A-36-16). Sites included three inshore (Punta Donato=PD, STRI Point=SP, Cristobal Island=CI) and three offshore (Bastimentos North=BN, Bastimentos South=BS, Cayo de Agua=CA) sites in the Bocas del Toro reef complex (BTRC), Panamá. Colonies were maintained in flow-through seawater at the Smithsonian Tropical Research Institute in BTRC prior to being shipped to the University of North Carolina at Chapel Hill overnight.

Upon return, small tissue samples (∼1 cm^2^) were collected from each colony and holobiont DNA was extracted (N=54) using a modified phenol-chloroform method as in (*53*). DNA extracts were cleaned using Zymo Genomic DNA Clean and Concentrator kits and concentrations were assessed using a Quant-iT PicoGreen dsDNA assay kit (Thermo Fisher). Samples of sufficient concentration (51/54 putative genotypes) were prepared for 2b-RAD sequencing (2b-RADseq; *54*), with 10 technical replicates to enable clone identification. A total of 61 samples were successfully sequenced across one lane of Illumina HiSeq 2500 using single-end 50 bp sequencing at the Tufts University Core Facility (TUCF).

2b-RADseq analysis followed the pipeline presented at https://github.com/z0on/2bRAD_denovo (accessed August 2021). Raw reads were trimmed and demultiplexed, cutadapt (*55*) removed reads with Phred quality score less than 15 and reads <36 bp in length. Because no *S. siderea* genome is available, a *de novo* reference was created. Following Rippe et al. (*33*), Symbiodiniaceae contamination was removed by mapping reads to concatenated genomes from four Symbiodiniaceae genera: *Symbiodinium* (*56*)*, Breviolum* (*57*)*, Cladocopium* (*58*), and *Durusdinium* (*59*) using Bowtie2 v2.4.2 (*60*). Putative symbiont contamination was removed and CD-HIT v4.7 (*61*) clustered and assembled remaining reads into a *de novo* reference consisting of 30 pseudochromosomes. Reads were mapped to this *de novo* reference using Bowtie2 v2.4.2 with default parameters and ANGSD v0.935 (*62*) was used for genotyping (using likelihood estimates) and identifying single nucleotide polymorphisms (SNPs). Standard filters were used to retain loci, which included ≥2x coverage, loci present at in at least 80% of individuals, a minimum mapping quality score of 20, a minimum quality score of 25, a strand bias p-value >1 x 10^-5^, a heterozygosity bias >1 x 10^-5^, a SNP p-value of 1 x 10^-5^, a minimum minor allele frequency >0.05, excluded all triallelic sites, and removed reads with multiple best hits. To distinguish putative clones, a hierarchical clustering tree (*hclust*) was constructed based on pairwise identity by state (IBS) values across all samples. Clones were determined using the similarity of technical replicates as a cut-off, and only one pair of clones was detected at PD (I4G and I4F; Fig. S1C). The clone pair with the lower total read count (I4G) was removed from all further analyses.

Population structure on the data set with the clone removed (8,105 SNPs) was determined using three methods: 1) hierarchical clustering of pairwise IBS values, 2) principal coordinate analysis (PCoA) based on IBS matrix, and 3) admixture proportions of individuals across sites. A height of 0.265 was used as the cut-off from the clustering dendrogram to distinguish three lineages (Fig. S1D). PCoA was performed on the covariance matrix in R v4.0.2 (*63*) using *capscale* (with null model, package=*vegan*, *64*), and was used in combination with the hierarchical clustering results to determine an optimal K of three. NgsAdmix v1.3.0 (*65*) with K=3 then determined the proportion of each individual’s ancestry that corresponded to each lineage (lineage 1 (L1), lineage 2 (L2), lineage 3 (L3)).

To estimate genetic differentiation, a different set of filters were used to retain loci, which included loci present in at least 80% of individuals, a minimum mapping quality score of 25, a minimum quality score of 30, a strand bias p-value >1 x 10^-5^, a heterozygosity bias >1 x 10^-5^, excluded all triallelic sites, removed reads with multiple best hits, and passed the lumped paralogs filter (770,398 SNPs). Genetic differentiation between lineages was estimated using ANGSD to find site allele frequency (SAF) for each lineage, after which realSFS determined the site frequency spectrum (SFS) for all lineage pairwise comparisons. Calculated SAFs and SFSs were then used to calculate global F_ST_, reported as weighted global F_ST_ values. Pearson’s Chi-squared test was used to determine if the distribution of L1 and L2 was dependent on reef type (inshore or offshore), excluding L3 individuals. All statistical analyses and data visualizations presented here were performed using R v4.0.2 (*63*).

### Characterizing baseline bacterial and Symbiodiniaceae communities

To characterize bacterial and Symbiodiniaceae communities associated with these *S. siderea* lineages, metabarcoding libraries were generated using a series of PCR amplifications for the V4/V5 region of the bacterial 16S rRNA gene (*66*, *67*) and the ITS2 region of Symbiodiniaceae ribosomal DNA (*68*, *69*), respectively. These libraries included both the colonies described above (but with L3 excluded; baseline time point) as well as fragments of these same colonies after a 50-day DTV experiment (post-DTV time point, described in detail below). Samples were sequenced (paired-end 250 bp) on an Illumina Miseq at TUCF and were analyzed together. For additional details on library preparation, refer to supplementary materials.

16S sequencing data were analyzed using *DADA2* (*70*), which conducted quality filtering and identified 17,903 amplicon sequence variants (ASVs) in 131 successfully sequenced samples (L3 individuals excluded). ASVs matching mitochondrial, chloroplast, and non-bacterial sequences were removed (1,487 ASVs) followed by an additional 160 ASVs identified in negative controls by the package *decontam*, leaving 16,256 ASVs and 131 samples with complete metadata remaining (N=47 baseline samples, N=84 post-DTV samples). Taxonomy was assigned using the Silva v132 database (*71*) and by using *blast+* against the NCBI nucleotide database (*72*). ASVs were checked for eukaryotic contamination, but none was detected. Counts were rarefied to 1000 reads per sample using the package *vegan*, leaving 6,144 ASVs and 47 samples for the baseline time point and 5,799 ASVs and 69 samples for the post-DTV time point. ASV richness and evenness in addition to Shannon and Simpson’s diversity indices were calculated from cleaned data (contaminant ASVs removed, but non-rarefied) using the *estimate_richness* function (package=*phyloseq*; *73*). An ANOVA tested for differences in diversity metrics across fixed effects of host lineage for baseline and host lineage plus DTV treatment for post-DTV. Bray-Curtis dissimilarity principal coordinate analyses (PCoAs) were conducted on all ASVs (relative abundance of rarefied datasets) using *phyloseq* for samples with complete lineage and DTV treatment metadata. *Vegan* and *pairwise.adonis* were implemented for statistical analyses of bacterial community differences and dispersion. 16S analyses were conducted on both rarefied and non-rarefied data, which resulted in similar patterns.

ITS2 data were submitted to SymPortal (*31*) to identify ITS2 type profiles. Successfully sequenced samples with corresponding metadata and excluding L3 individuals (N=47 baseline samples, N=84 post-DTV samples) were visualized for differences between host lineages for baseline samples, and host lineages and DTV treatment for post-DTV samples. Defining intragenomic variants (DIVs) were summed by majority ITS2 sequence before calculating relative abundances. Relative abundances of majority ITS2 sequences were compared with bar plots using *phyloseq*, and a Kruskal-Wallis test (*kruskal.test*) tested for differences in the proportion of D1 majority ITS2 sequences across L1 and L2.

### Assessing baseline holobiont phenomes and skeleton morphologies

Coral colonies were sectioned into fragments using a tile saw (RIDGID) and affixed to pre-labeled plastic petri dishes using cyanoacrylate glue (IC-Gel Insta Cure; BSI). All fragments were maintained in aquaria for 16 days at 28°C with standardized light conditions of 400 µmol photon m^2^ s^-1^ on a 12 h:12 h light:dark cycle using full spectrum LED lights (Euphotica; 120W, 20000K) based on Rodas et al. (*74*). After this recovery period, baseline phenomic metrics of coral host and algal symbiont health were assessed by flash freezing one fragment of each genotype. Flash frozen fragments were thawed and tissue was removed via airbrush and seawater, homogenized, and centrifuged to separate host and symbiont fractions, which were divided into four aliquots. One symbiont aliquot was used for symbiont cell counts and all other host and symbiont aliquots were disrupted using a bead mill homogenizer (Omni Bead Mill 24; GA, USA) with a high throughput hub at 6 m s^-1^ for 2 min for downstream phenomic assessments.

Total host protein was quantified using the Bradford method (*75*) with absorbances read at 595 nm on a microplate reader (Biotek Synergy H1; CA, USA). Data were converted from absorbance to total protein concentrations (μg μL^-1^) using a standard curve of Bovine Albumin Serum and then normalized to surface area. Total host and algal symbiont carbohydrates were quantified using the phenol-sulfuric acid method (*76*) with absorbances read at 500 nm. Carbohydrate values (mg mL^-1^) were calculated from raw absorbance using a D-glucose standard curve and then normalized to surface area. This method measures all monosaccharides, which includes glucose, the main byproduct of photosynthesis that is translocated from algal symbiont to coral (*77*). Algal cell density was quantified in triplicate using the hemocytometer method (*78*) and then normalized to surface area. Symbiont photosynthetic pigments (chlorophyll *a* = Chl *a*) were measured spectrophotometrically, read at 663 nm and 630 nm, calculated following equation 1 below (*79*) where *A663* represents blank-corrected absorbance values at 663 nm and *A630* represents blank-corrected absorbance values at 630 nm, and then normalized to surface area.

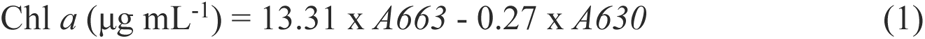

Tissue thickness (mm) for all fragments was measured with calipers after tissue was removed and corallite surface area (mm^2^) was measured followed methods presented by Conti-Jerpe et al. (*80*). Briefly, the polygon tool in ImageJ measured the area of seven corallites from each fragment in pixels, which was converted to mm^2^ using a size standard in each photograph.

Skeletons from the baseline time point were scanned at high resolution (>1200 dpi; Epson Perfection V550) and additional skeleton morphometrics were quantified using ImageJ. Measurements were taken randomly from 5-10 corallites per fragment that were perpendicular to the field of view and not in a budding state nor at the colony margins to avoid areas of recent growth. Several skeletal morphometric traits were measured, including: calyx diameter, septal length and width from the primary cycle, inter-corallite (*i.e.,* theca) wall thickness, corallite spacing, and corallite density (# corallites cm^-2^).

Light enhancement factor (LEF), which describes the optical property of the skeleton to collect and enhance the light field, was estimated by quantifying relative changes in the intensification of local illumination on a naked coral skeleton due to multiple light scattering (*81*). Before measuring LEF, coral skeletons were soaked in 10% commercial bleach for 48 h and dried at 50–60°C for 48 h to avoid absorption from remaining organic matter. Detailed descriptions of the method can be found in (*28*), but briefly LEF was measured in triplicate for each fragment and expressed as the ratio of reflected light from a skeleton illuminated by a light source and the reflected light from a reference (black fabric). Illumination was provided by a linearly polarized continuous wave laser diode (532 nm) coupled with a lens to a miniature isotropic probe fabricated with standard fused silica step-index optical fibers (numerical aperture=0.22, 200 μm diameter core). This probe was used as a light source and placed in proximity to the cavities of the skeleton. The light scattered by the skeleton was measured using a miniature spectroradiometer (Ocean Optics, Inc.). While LEF reflects the maximum potential ability of the coral skeleton to enhance the light field for algal photosynthesis in the coral tissue (*82*), it is not a descriptor of coral tissue pigmentation.

To characterize baseline holobiont phenomic differences across lineages, all host and symbiont physiology metrics were *log*-transformed and combined in principal component analyses (PCAs) using the *FactoMineR* package. Significance of the fixed effect of lineage was assessed with PERMANOVA, using the *adonis* function (package=*vegan*). Effect size was determined using partial Omega-squared (𝜔^2^) values calculated with the *adonis_OmegaSq* function (package=*MicEco*). The same method and PERMANOVA model that were used to assess baseline phenome PCAs was also used for the skeleton morphology PCA. A separate linear model was used to assess the main effect of lineage on light enhancement factor (LEF). For all linear models presented here, assumptions and model fit were assessed visually using the *check_model* function (package=*performance*) and pairwise comparisons were calculated using *emmeans*.

### Testing responses to thermal variability and thermal challenge across lineages

To determine how host lineages respond to thermal variability and thermal challenge, we exposed fragments of each colony to a 50-day diel thermal variability (DTV) experiment, followed by thermal challenge and recovery. *In situ t*emperature data were continuously collected every 15 minutes for approximately one year prior to colony collection, with HOBO ProV2 temperature loggers (Onset, Bourne, MA) deployed between 1 and 4 m depth at each of the six sites (Fig. S2A) from June 10, 2015 to August 14, 2016. Loggers were recovered from four sites, including three inshore (CI, PD, SP), and one offshore (CA). Differences in daily mean temperature and daily temperature range (reported in Table S1A) across sites were determined using a one-way ANOVA with Tukey’s HSD post-hoc tests (Fig. S2B; Table S13). Two DTV treatments were designed based on these *in situ* temperature data. The control treatment (no DTV) was maintained at 29.5℃, representing the overall daily mean of all sites (29.6 ± 0.02℃; Table S1). The DTV treatment had a minimum temperature of 28.5℃ with daily increases of 3°C (28.5 - 31.5℃) and this treatment was informed by the highest *in situ* DTV observed (3.2℃ at CI) (Fig. S2, Table S1).

Colony fragments (N=18 genotypes per tank, 3 aquaria per treatment) were randomly distributed into treatment aquaria for 16 days at 28°C (recovery), followed by 15 days of acclimation to experimental conditions. Next, a 50-day DTV experiment was conducted followed by a 15-day thermal challenge (32°C) and 16-day recovery period (Fig. S3C). Differences in daily temperature parameters (variability, mean, maximum, and minimum) across treatments during the 50-day DTV treatment were determined using a one-way ANOVA, and the effect size (*i.e.,* variance in temperature parameter explained by experimental treatment) was determined using Eta-squared (*η^2^*) values calculated using the *eta_squared* function (package=*effectsize*). Distinct treatments were maintained over the course of the 50-day DTV treatment (Fig. S10; Table S14). Throughout the experiment, light conditions were maintained at 400 µmol photon m^2^ s^-1^ on a 12 h:12 h light:dark cycle. Corals were fed freshly hatched *Artemia sp.* nauplii two to three times weekly, at night, and were allowed to feed for one hour before resuming recirculating flow in the aquaria. For more detailed experimental information, including water quality, see supplementary materials.

Coral growth was estimated using the buoyant weight method under standard conditions (28°C and 33 ppt) with a bottom-loading balance (precision=0.0001 g; Mettler-Toledo, Columbus, OH) at four time points: after acclimation (T_0_), during DTV (T_1_), at the end of DTV (T_2_), and at the end of recovery (T_3_) (see experimental timeline, Fig. S2C). Growth was calculated as a specific growth rate per day through DTV treatment (T_0_ to T_2_) as well as through heat stress and recovery (T_2_ to T_3_). Immediately after buoyant weighing, corals were imaged to quantify surface area for physiology standardization with a CoralWatch Health Chart (*83*). Distance between the camera and corals as well as lighting were standardized. Surface area measurements were obtained using ImageJ (*84*) and only live tissue was included in surface area normalizations. The effect of DTV and host lineage on growth was assessed separately for two durations: 1) throughout the 50-day DTV treatment and 2) during the thermal challenge and recovery periods. For both durations, linear mixed models (package=*lme4*) were implemented with main effects of treatment and lineage and a random effect of genotype.

Photochemical efficiency of photosystem II (Fv/Fm) was measured in triplicate for each fragment using a Diving PAM (Walz) at seven time points: once at the end of DTV, three times during thermal challenge, and three times during recovery. For all time points, corals were dark-acclimated for at least 30 minutes before measurements, which started between 17:00 and 19:00 and ended between 20:00 and 22:00. Measurements were made using a saturation pulse width of 0.6 s at full strength light intensity, electronic signal damping of 2, and gain of 4. A linear mixed model was implemented to assess the interactive effects of time, lineage, and DTV treatment (with a random effect of coral genotype) on Fv/Fm throughout the thermal challenge and recovery periods.

Phenomic metrics of holobiont health were assessed by flash freezing fragments following final recovery and following the same methods used for baseline holobiont phenomes described above. To characterize phenome-wide responses, all metrics were *log*-transformed and combined in PCAs using the *FactoMineR* package. Significance of each factor (fixed effects of DTV treatment and host lineage) were assessed with PERMANOVAs, and effect size of each factor was determined using 𝜔^2^ values calculated as described above. For all end of experiment models (*i.e.,* when more than one factor of interest was present), models were selected based on a backwards selection method, where only significant interaction terms were maintained in the model.

Coral holobiont DNA was also subsampled, preserved in 100% ethanol and stored at −80°C at the end of the 50-day DTV treatment to assess changes in bacterial and Symbiodiniaceae communities in response to DTV, following methods described above (Fig. S6 and S7, respectively).

### Historical growth estimates via sclerochronology

In 2015, a total of 39 cores of large *Siderastrea siderea* (∼1 m diameter) colonies were collected from four pairs of inshore-offshore reef sites in BTRC (permit SE/A-28-15) using a hydraulic core drill following methods previously described in detail (*49*, *85*). The eight sites where cores were collected include the six sites detailed above (Fig S2A; PD, SP, CI, BN, BS, CA), as well as two additional sites (inshore: Punta Laurel=PL; offshore: Drago Mar=DM). From each site, five healthy colonies were randomly selected for coring. In some cases, less than five colonies were sampled due to a lack of corals of sufficient size. Skeletal cores were preserved in 200-proof ethanol and transported to UNC-Chapel Hill, where they were scanned using X-ray computed tomography to visualize annual density growth bands (following *49*, *85*). In brief, boundaries between semiannual density bands were manually delineated, and linear transects were traced down the central axis of three corallites to estimate growth (Fig. S9B). Following a previously established protocol (*86*), nine density standards were included in each scan to facilitate converting skeletal density measurements from CT Hounsfield units to g cm^-3^. Linear extension was measured in the image viewing software Horos v2.0.2 as the width of each annual density band couplet, and calcification (g cm^-2^) was calculated as the product of skeletal density and linear extension.

In 2023, tissue was scraped from the surface of remaining *S. siderea* cores (N=38), and DNA was isolated using Qiagen Blood and Tissue kits following manufacturer’s instructions. DNA extracts were cleaned and prepared for 2b-RADseq following methods detailed above. Samples were sequenced at TUCF along with replicate samples from the original lineage dataset to facilitate identifying lineage of each core. After lineage assignment, N=24 cores remained with both growth chronology and lineage assignment. A linear model with fixed effect of lineage was used to test for lineage differences in skeletal density, linear extension, and calcification separately for two separate time intervals: 1) all data (1880–2014) and 2) recent data (1980–2014). Analyses was separated into these two time intervals to confirm lineage-specific historical growth patterns were consistent regardless of the number of data points per core (*i.e.,* core age), as sample size decreases moving back in time. The linear model results were consistent regardless of time interval; therefore, results from analyses of recent data are presented in the main text and results from all data are found in the supplemental material.

Monthly historic sea surface temperature (SST) data were obtained from 1870 to 2023 from the Hadley Centre Sea Ice and Sea Surface Temperature (HadISST) dataset (*87*) for the 1° latitude-longitude grid that covered all sampling sites. The HadISST database combines *in situ* and satellite observations of global SST data, making it an ideal resource for assessing historic trends in temperature (*e.g., 88*, *89*). Monthly temperature observations were averaged to estimate annual mean (January-December) and annual summer mean (August-November) for the region. Linear models with fixed effects of temperature (either annual summer mean or annual mean) and lineage were used to explore whether historical calcification was modulated by temperature changes and whether these changes were different between lineages. These linear models were completed with data from the two time intervals of historical growth data (*i.e.,* all data and recent data). Additionally, a separate linear model was used to assess whether annual mean and annual summer mean increased between 1870 to 2023 in BTRC.

## Acknowledgments

We extend appreciation to the staff at the Smithsonian Tropical Research Institute at Bocas del Toro, especially Urania Gonzalez for assistance coordinating field work. Thanks to our fearless boat captain Sebastian Castillo, in addition to Clare Fieseler and Logan Buie for field assistance. Thank you to Jack Weldon for assistance with physiology assays. Analysis was made possible through BU’s Shared Computing Cluster.

## Author contributions

Conceptualization: SWD, KDC, HEA

Methodology: PG, BEB, SWD, HEA, JHB, JPR, KDC

Investigation: HEA, BEB, KGC, MIMR, LT, OCN, DS, AMP, NGK, JEF, CBB, AMH

Visualization: HEA

Supervision: HEA, SWD, KDC

Writing—original draft: HEA, SWD

Writing—review & editing: HEA, BEB, KGC, MIMR, JEF, LT, AMH, CBB, OCN, AMP, DS, NGK, JHB, JPR, PG, KDC, SWD

## Data and materials availability

Sequence reads will be uploaded to NCBI’s sequence read archive (SRA) upon acceptance. All other data, code, and materials used in the analyses can be found on the Github repository associated with this project: https://github.com/hannahaichelman/CrypticCorals

