## Supplemental Materials for "Cryptic diversity shapes coral symbioses, physiology, and response to thermal challenge"

Hannah E. Aichelman *et al.*

**Table of Contents for Supplementary Materials:**

- Supplementary Results
- Supplementary Materials and Methods
- Figs. S1 to S10
- Tables S1 to S14

### Supplementary Results

#### *Aquaria conditions during DTV treatment*

Distinct treatments were maintained during the 50-day DTV treatment (Figure S10), and mean DTV was significantly higher in DTV aquaria relative to controls ($p$<0.001; Figure S10A). While daily mean temperatures were significantly higher in control relative to DTV aquaria ($p$=0.04; Figure S10B), the effect size of this difference was marginal ($\eta^2$=0.04) relative to the much stronger effect size of DTV ($\eta^2$=0.97; Figure S10A).

#### *Aquaria conditions during thermal challenge and recovery*

Following the end of the DTV treatment, temperatures in all aquaria were brought to the thermal minimum and then increased 1°C per day to an 8-day heat challenge treatment with mean (± standard error) temperature across the two treatments of 31.8°C (± 0.007°C). Subsequently, temperatures were decreased by 1°C per day to recovery conditions, which were maintained for 16 days at a mean (± standard error) of 29.3°C (± 0.008°C) (Fig. S2C). Following the recovery period, all coral fragments were frozen.

### Supplementary Materials and Methods

#### *Experimental conditions*

Throughout the experiment, corals were maintained in 42 L aquaria at salinity of 33 ppt using Instant Ocean Sea Salt artificial seawater (ASW), which was refreshed weekly with 40% water changes. Nubbins were rotated weekly to ensure even light exposure over the course of the experiment. Salinity was measured in each treatment at least once daily using a YSI 3200 conductivity meter (Yellow Springs, Ohio, USA), and pH was measured 3 times weekly in each treatment using an Orion Star A211 pH meter (calibrated with certified NBS pH buffers of 4.01, 7.00, and 10.01). Temperature was controlled with two 500 W heaters (Eheim) and a chiller (AquaEuroUSA) that were used to maintain DTV profiles in each treatment using the Neptune Systems Apex Classic AquaController with custom coded profiles. Briefly, virtual heating and cooling segments were used to dictate changes in target temperature for each treatment in two-hour intervals and the DTV treatment descended to a nighttime minimum of 28.5°C for 9 hours. Temperature was measured daily using a NIST-calibrated glass thermometer and evaluated against Apex readouts, with twice weekly calibration. Temperature was recorded by the Apex system and at 5-minute intervals with HOBO ProV2 Loggers (Onset, Bourne, MA) throughout the duration of the experiment, except for the DTV treatment, which is missing data for the heat challenge and recovery periods due to a failed logger. Therefore, HOBO logger data during the heat challenge and recovery periods was supplemented with the semi-regular readings from the NIST-certified glass thermometer (Fig. S2C). Experimental temperature data presented here is HOBO logger data corrected to the calibrated Apex data, as this is the most complete temperature record across the treatments.

***Profiling of prokaryotic and Symbiodiniaceae communities***

ITS2 primers (*68, 69*) included the forward SYM_VAR_5.8S2 (5’ - TCGTCGGCAGCGTC AGATGTGTATAAGAGACAG *NNNN* **GTGAATTGCAGAACTCCGTG** - 3’) and the reverse SYM_VAR_REV (5’ - GTCTCGTGGGCTCGG AGATGTGTATAAGAGACAG *NNNN* **CCTCCGCTTACTTATAGCTT** 3’). 16S primers (*66, 67*) included the forward Hyb515F (5’-TCGTCGGCAGCGTC AGATGTGTATAAGAGACAG *NNNN* **GTGYCAGCMGCCGCGGTAA**-3′) and the reverse Hyb806R (5’-GTCTCGTGGGCTCGG

AGATGTGTATAAGAGACAG *NNNN* **GGACTACNVGGGTWTCTAAT**-3’). Underlined bases denote adapter linker, bold bases are primer sequences, and the middle bases are spacer sequences. Both ITS2 and 16S PCR reactions totaled 30 µl and included 30 ng of template DNA, 1 µM forward primer, 1 µM reverse primer, 0.2 mM dNTP, 1X ExTaq buffer (Takara), 0.025 U ExTaq enzyme (Takara), and the remaining Milli-Q $H_2O$ (Millipore). For ITS2 primers, the reaction profile cycled at 95°C for 40 seconds, 59°C for 120 seconds, and 72°C for 60 seconds for 28 cycles and a final elongation step of 72°C for 7 minutes. The reaction profile for 16S primers cycled at 95°C for 40 seconds, 58°C for 120 seconds, and 72°C for 60 seconds for 32 cycles with a final elongation step of 72°C for 5 minutes. For 16S, two negative controls using water were prepared and later used to remove contaminant sequences. PCR products were purified using GeneJET PCR Purification kits (ThermoFisher) eluted in 30 µl. Each PCR product was uniquely barcoded, subjected to five PCR cycles, and visualized on a 1% agarose gel to assess relative concentrations. Samples were pooled in equal concentrations within libraries, and 25 µl of the pooled library was run on a 1% SYBR Green (Invitrogen) stained gel. The target band was excised and incubated with 30 µl of Milli-Q water overnight at 4˚C. The ITS2 and 16S libraries were then quantified using a Quant-iT PicoGreen dsDNA assay kit (Thermo Fisher), pooled based on concentration (1:3 ITS2 and 2:3 16S) and submitted for paired-end 250bp sequencing on an Illumina Miseq at TUCF.

***Supplemental statistical analyses***

The effect of lineage on individual host and symbiont physiology metrics (*i.e.,* total protein, total host and symbiont carbohydrate, chlorophyll *a*, and symbiont density) were assessed separately for two time points: 1) at baseline, and 2) at the end of the experiment. To assess physiology data collected at baseline, a linear model was implemented with a main effect of lineage, and for the end of the experiment a linear mixed model was implemented with main effects of lineage and DTV treatment (with a random effect of genotype). Additional linear models were used to assess the main effect of lineage on corallite area and light enhancement factor (LEF) (Figure S5; Table S7).

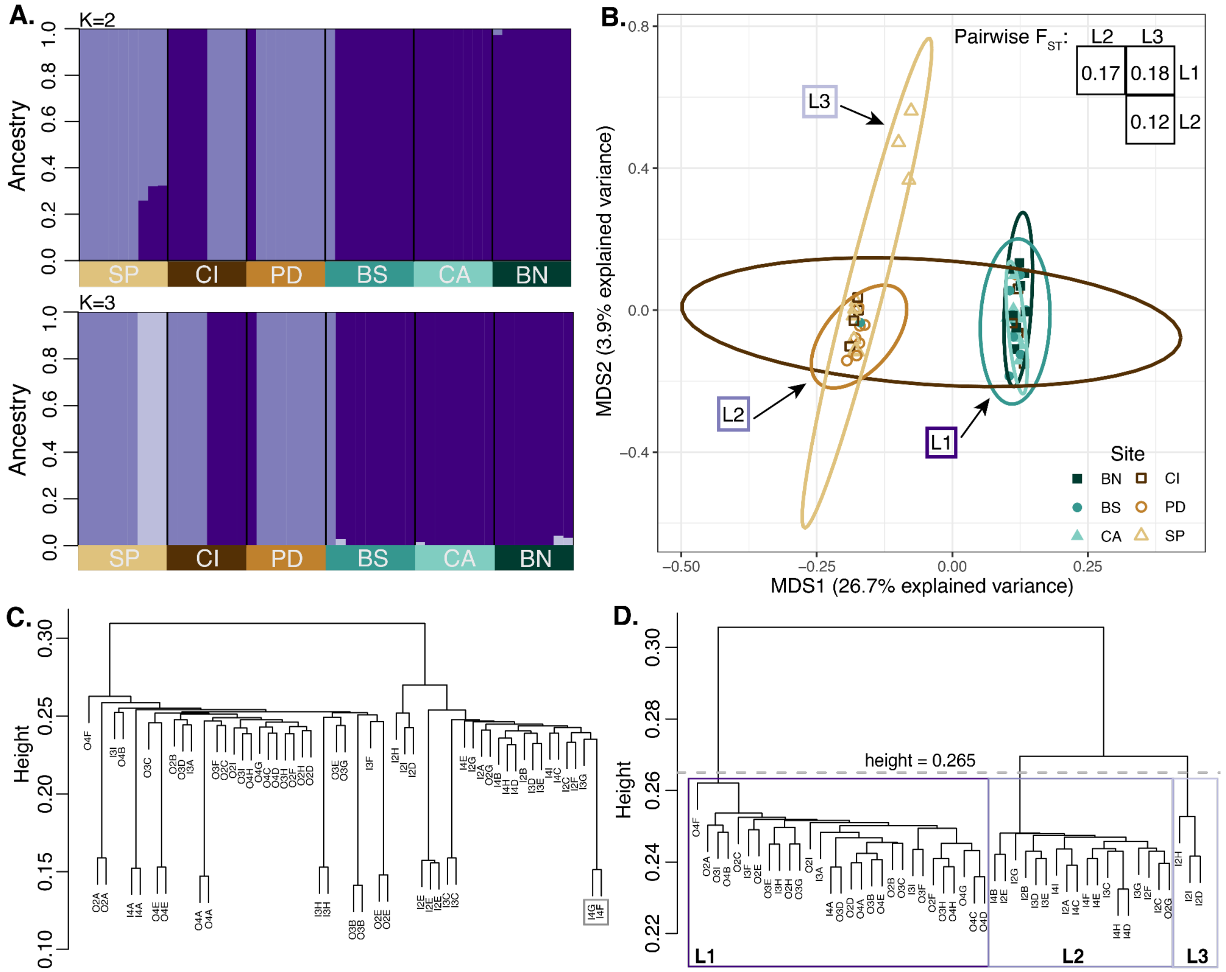


**Figure S1. 2bRADseq identifies three cryptic lineages of a *Siderastrea siderea* species complex in the Bocas del Toro Reef Complex, Panamá.**

**A.** ADMIXTURE results for K=2 (top) and K=3 (bottom). Columns represent an individual and the bar color represents an individual's assignment to one of two (top) or three (bottom) ancestral populations (L1=dark purple, L2=medium purple, L3=light purple). Boxes below group individuals by collection site (colors as in B). **B.** Principal coordinate analysis (PCoA) plot illustrating clustering of lineages across collection sites (N=50 individuals). Shapes and colors distinguish sites, with closed shapes/green shades representing offshore sites and open shapes/brown shades representing inshore sites. Inset shows pairwise $F_{ST}$ values between lineages. **C.** Identity by state (IBS) cluster dendrogram of all samples. The gray box highlights one putative clonal pair (I4G and I4F), which occurs at the same height as the technical replicates (indicated by identical sample name). **D.** IBS cluster dendrogram with technical replicates and one putative clonemate removed (I4G). The dashed line at height=0.265 represents the cutoff for lineage assignment, with lineages 1-3 (L1, L2, L3) indicated with boxes colored as in A. All L3 data were removed from downstream analyses due to low sample size.

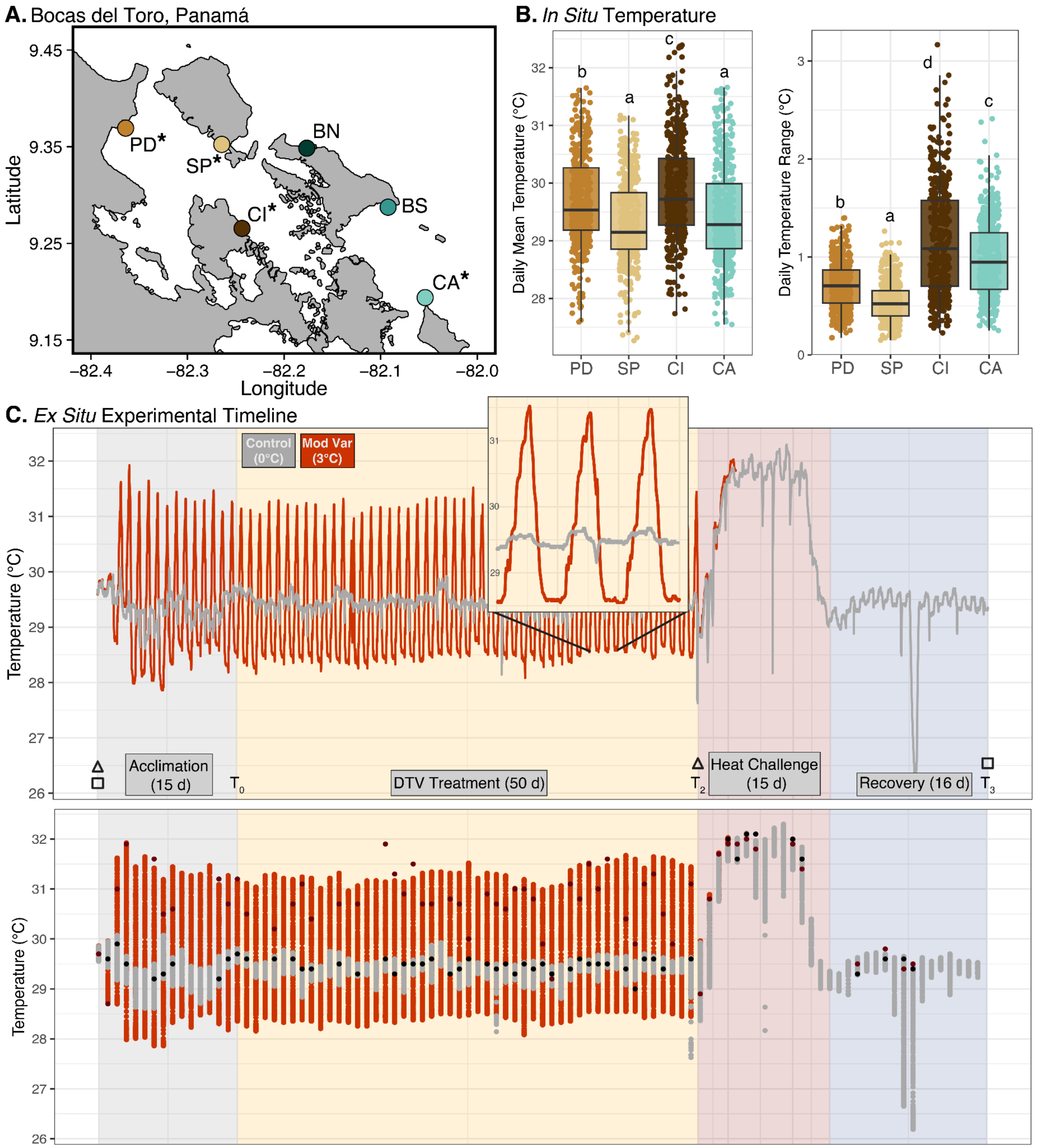


**Figure S2. Thermal variability experimental design.**

**A.** Map indicating six sites across the Bocas del Toro Reef Complex in Panamá where *Siderastrea siderea* colonies were collected, including three inshore (brown shades: PD=Punta Donato, SP=STRI Point, CI=Cristobal Island) and three offshore (green shades: BN=Bastimentos North, BS=Bastimentos South, CA=Cayo de Agua) sites. Asterisks indicate sites where temperature loggers were recovered. **B.** Daily mean temperature (left) and daily temperature range (right) recorded using HOBO loggers for approximately one year prior to coral collection (6/10/2015-8/14/2016). Distinct letters indicate significant differences in temperature parameters from ANOVA and Tukey's HSD post-hoc tests (Table S1; N=431 for each site). **C.** Timeline of *ex situ* diel thermal variability (DTV) experiment, including 15 days of acclimation, 50 days of DTV treatment, 15 days of heat challenge, and 16 days of recovery. *Ex situ* temperature data were obtained from HOBO loggers, which recorded temperature every 5 minutes, and was supplemented with semi-regular readings from a NIST-certified glass thermometer. The top panel indicates HOBO logger data only. Because the HOBO logger in the DTV treatment failed during the heat challenge, the bottom panel represents the same HOBO logger data in the top panel along with readings from the glass thermometer (black points=control, dark red points=DTV treatment). Squares indicate when fragments were flash frozen to measure coral and symbiont phenomic metrics. Triangles indicate when fragments were subsampled for DNA, at baseline to determine host genetics and post-DTV to assess Symbiodiniaceae and bacterial communities. $T_0$, $T_2$, $T_3$ indicate when corals were buoyant weighed to calculate growth. The inset in the top panel illustrates temperatures over three days to highlight DTV treatments in more detail.

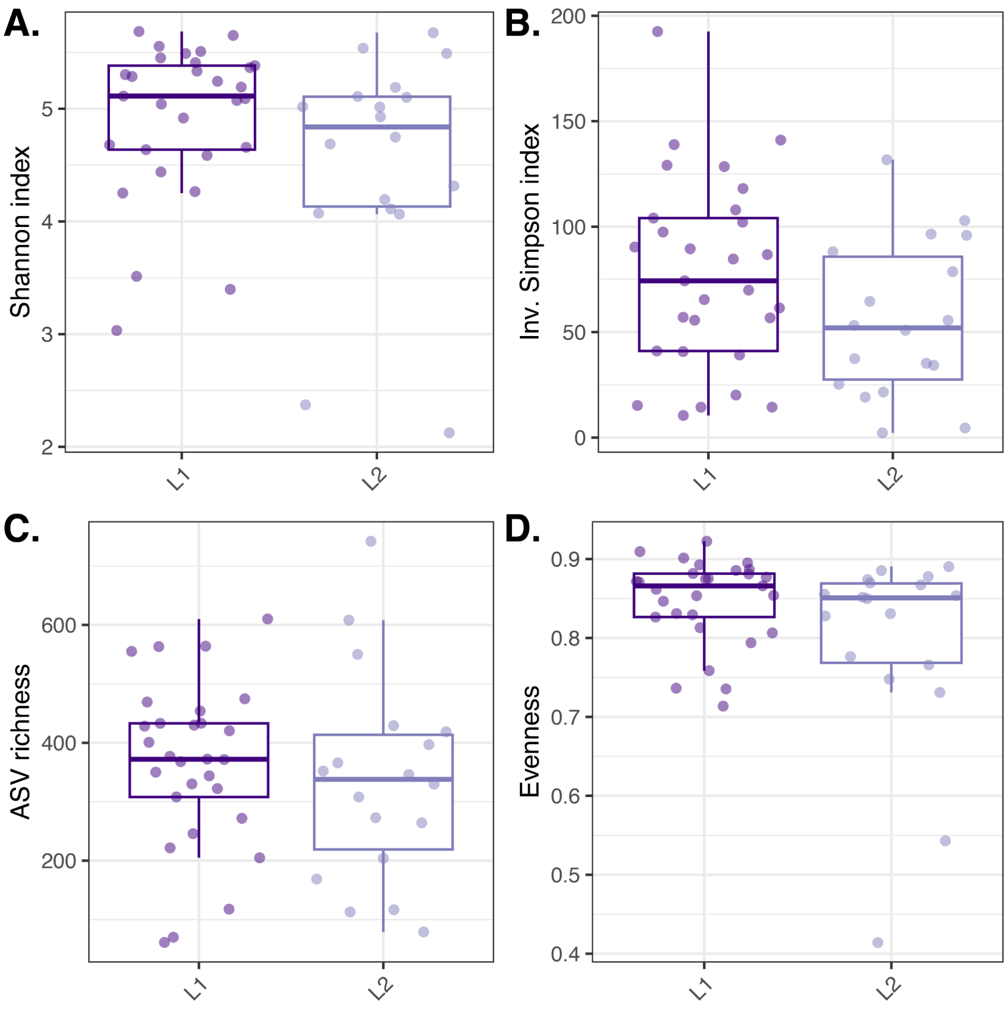


**Figure S3. Bacterial community diversity is not influenced by cryptic host lineage at baseline.**

Diversity metrics, including Shannon index **(A)**, Inverse Simpson index **(B)**, ASV richness **(C)**, and Evenness **(D),** presented by cryptic host lineage. All diversity metrics were calculated using cleaned data (contaminant ASVs removed, but not trimmed or rarefied) (N=47).

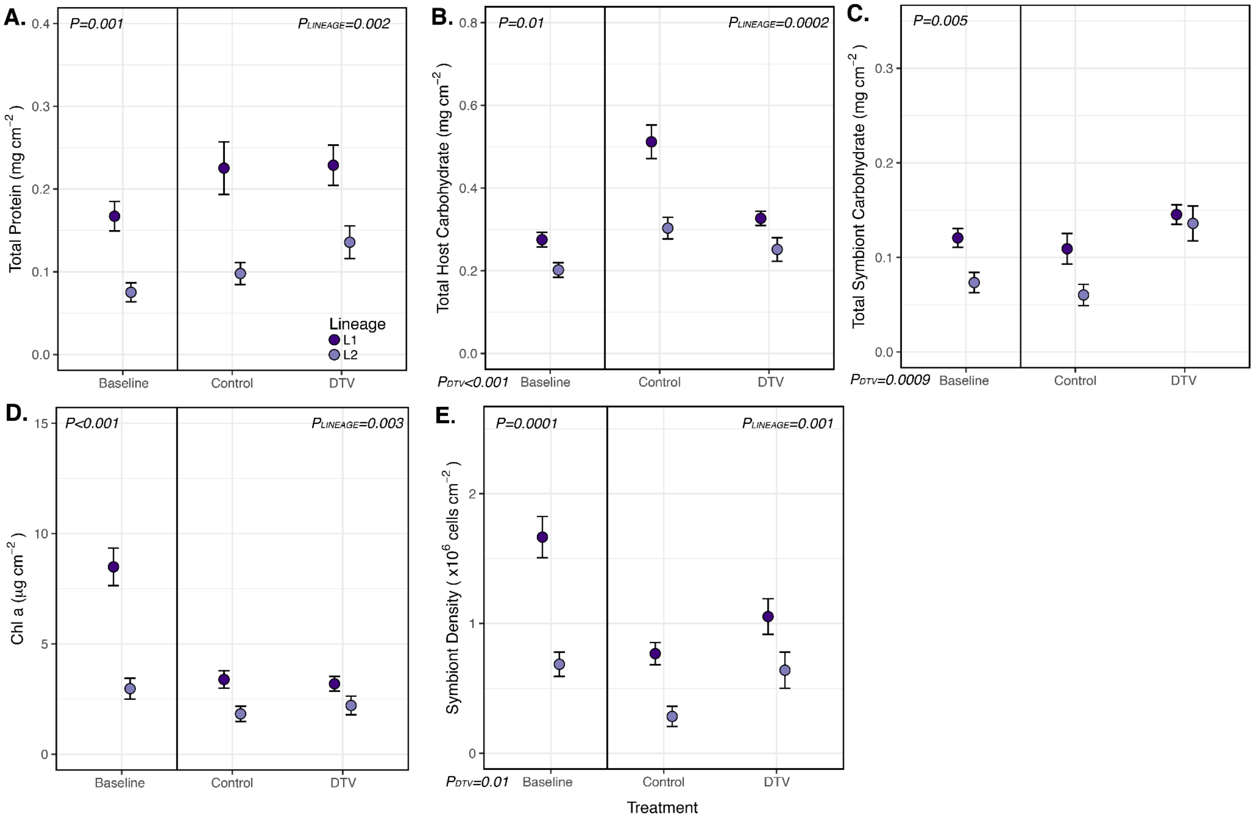


**Figure S4. Cryptic lineage differences in individual host and symbiont physiology metrics.** Individual physiology metrics (y-axis) measured throughout the experimental interval (x-axis), including at baseline (Treatment=Baseline) and at the end of the experiment (Treatment=Control or DTV). Points represent mean ± standard error of total protein (mg cm$^{-2}$; **A**), total host carbohydrate (mg cm$^{-2}$; **B**), total symbiont carbohydrate (mg cm$^{-2}$; **C**), chlorophyll *a* (µg cm$^{-2}$; **D**), and symbiont density (cells cm$^{-2}$; **E**). Statistical significance of lineage was assessed separately for each time point (*i.e.,* baseline or end of experiment), and significance is indicated on the plots within each time point. For example, a significant effect of cryptic host lineage on total protein values at baseline is indicated above the 'Baseline' data points in panel A. Statistical significance of DTV itself on the data from the end of experiment time point is indicated at the bottom-left of each panel.

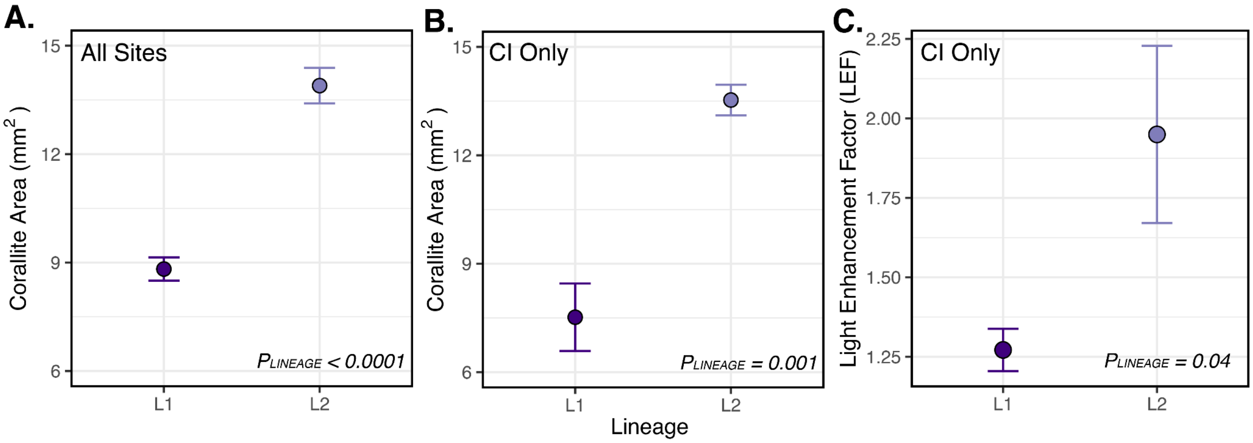


**Figure S5. Distinct skeletal morphology between cryptic host lineages.**
**A.** Mean corallite area ($mm^2$) ± standard error from corals from all collection sites showcasing that lineage 1 (L1; N=29) had smaller mean corallite area than lineage 2 (L2; N=16). **B.** The same differences in mean corallite area ± standard error were observed when considering L1 (N=4) and L2 (N=4) that co-occur at Cristobal Island (CI). **C.** Light enhancement factor (LEF) data presented in Figure 2D, except only considering when L1 (N=4) and L2 (N=3) co-occur at CI. P-values indicate statistical differences in corallite area between lineages according to linear models.

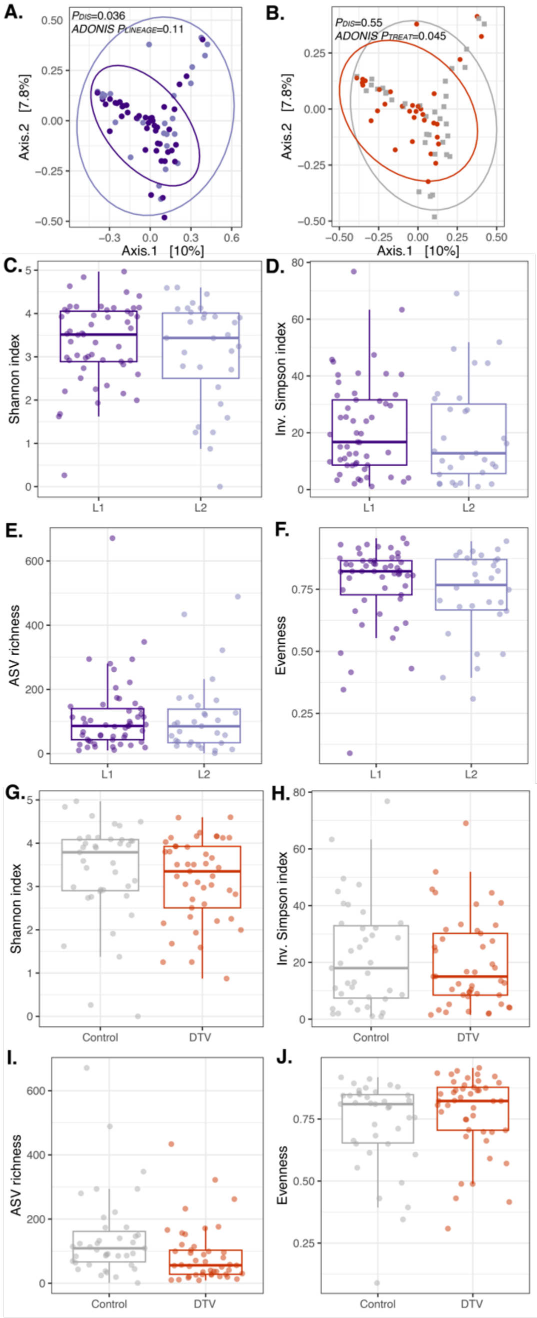


**Figure S6. Bacterial community diversity and structure after diel thermal variability (DTV) treatment (post-DTV).**

**A,B.** Bray-Curtis dissimilarity principal coordinate analysis (PCoA) of coral bacterial communities for all ASVs colored by lineage (**A**) and DTV treatment (**B**). Ellipses represent 95% confidence intervals, *ADONIS P*-values indicate significant differences in community beta diversity, and $P_{DIS}$ values compare dispersion across sites. PCoAs were calculated using cleaned and rarefied data (N=69). Diversity metrics, including Shannon index **(C,G)**, Inverse Simpson index **(D,H)**, ASV richness **(E,I)**, and Evenness **(F,J),** presented by cryptic host lineage **(A-D)** and DTV treatment (**E-H**). All diversity metrics were calculated using cleaned data (contaminant ASVs removed, but not trimmed or rarefied) (N=84).

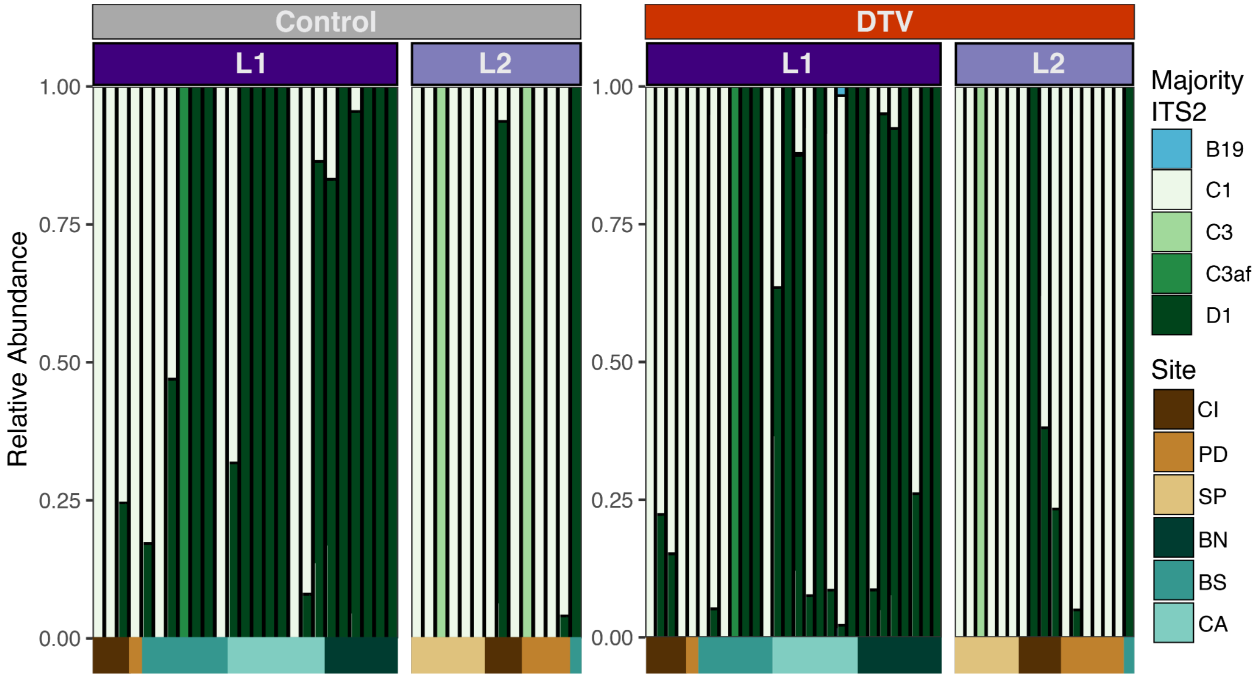


**Figure S7. Symbiodiniaceae communities are driven by cryptic host lineage, not diel thermal variability (DTV).**
Bar plots of Symbiodiniaceae majority ITS2 sequence relative abundance data, colored by majority ITS2 type. Defining intragenomic variants (DIVs) were summed by majority ITS2 sequence before calculating relative abundances. Bar plots are faceted by lineage (L1=dark purple, L2=light purple) within experimental treatment (Control and DTV). Each column of the bar plots represents a coral individual, and color blocks under the plots represent the site of origin for each individual.

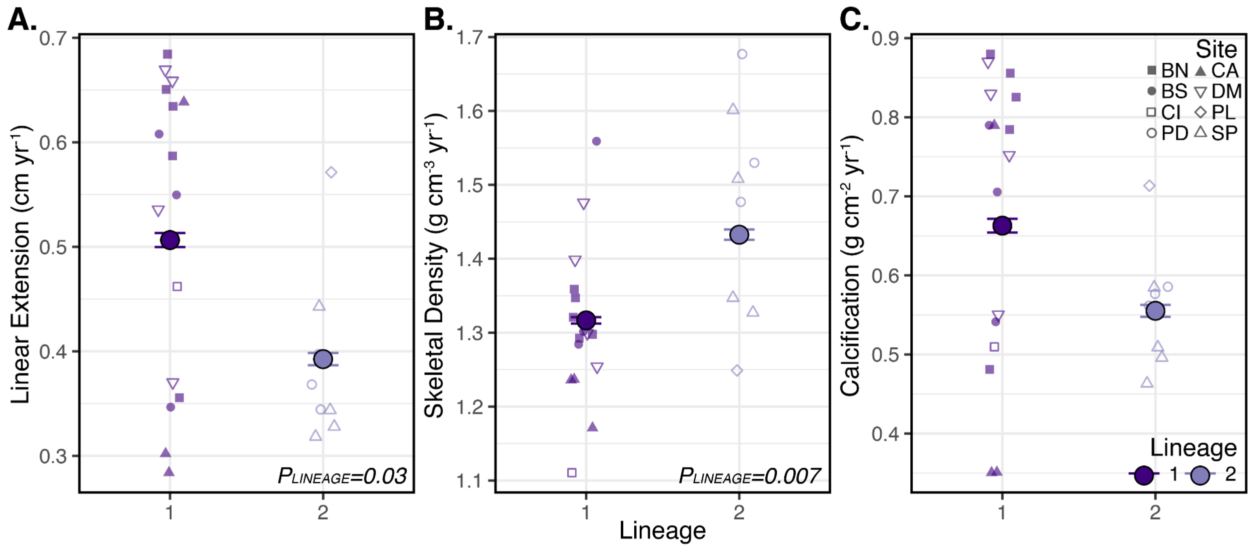


**Figure S8. Cryptic host lineage differences in historical growth including all growth records.**

Linear extension (**A**), skeletal density (**B**), and calcification (**C**) across cryptic host lineages. For all panels, large points represent the mean ± standard error of each metric for each lineage, and smaller points represent an individual coral core. Shape of the smaller points represents the site of origin for that core. The entire growth records of each core are included here (1880-2014; N=16 L1 cores, N=8 L2 cores).

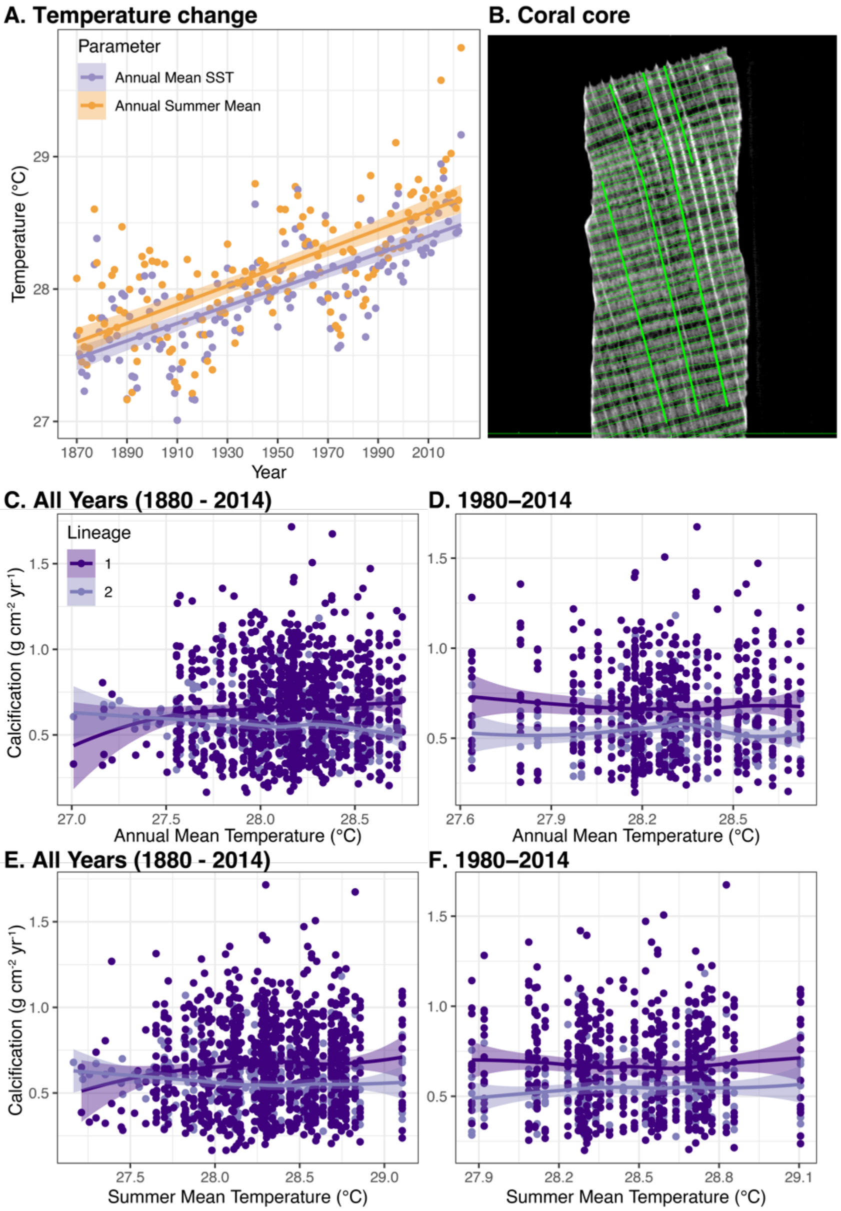


**Figure S9. Historical coral calcification rates are not related to temperature.** **A.** Annual mean sea surface temperature (SST) and annual summer mean temperature increased between 1870 - 2023 in the Bocas del Toro archipelago (both $p<0.001$). Lines in **A** represent the linear model fit and shaded areas indicate 95% confidence intervals produced by *geom_smooth()*. **B.** Example of measuring linear extension with the image viewing software Horos v2.0.2. **(C-F).** There was no relationship between historical calcification and annual mean temperature (**C**,**D**) or summer mean temperature (**E**,**F**) across cryptic host lineages for any time interval (all data=1880-2014=**C**,**E**; recent data=1980-2014=**D**,**F**). Each point represents one year of calcification from one core at the representative temperature. Colors represent cryptic coral lineage, and sample size is the same as Figure S8 (N=16 L1 cores, N=8 L2 cores). Lines in **C-F** are fit using the loess function and shaded areas indicate 95% confidence intervals produced by *geom_smooth()*.

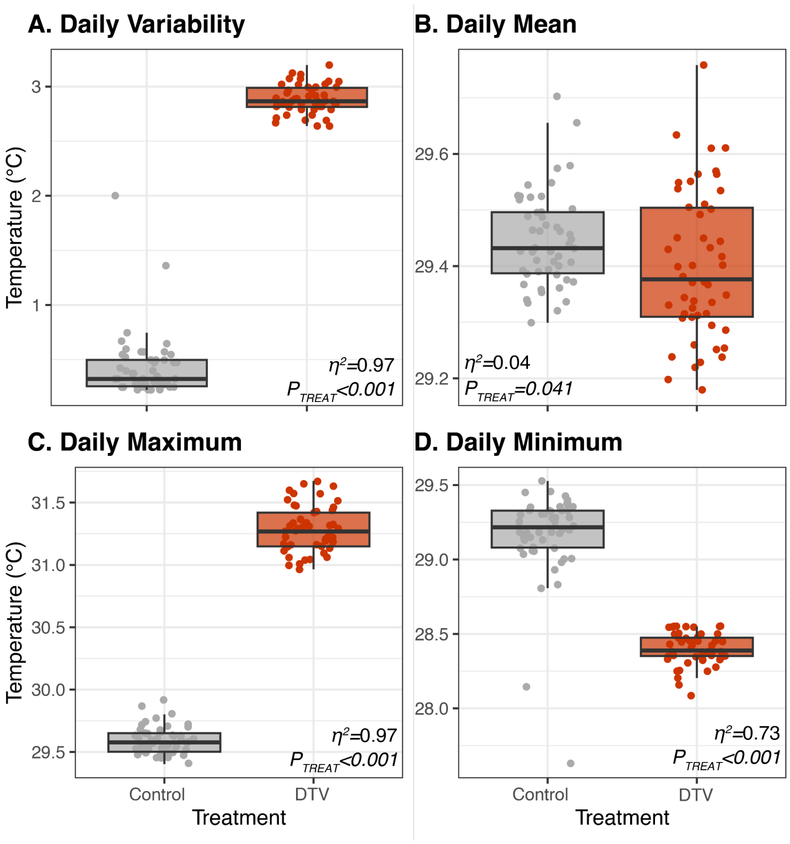


**Figure S10. Temperature characteristics of diel thermal variability (DTV) experimental treatments.**

Temperature data recorded using HOBO loggers for the duration of the 50-day DTV experiment (September 22, 2016 - November 10, 2016). For all box plots, each point represents one day of temperature data per treatment. Temperature metrics include: **(A)** diel thermal variability (DTV), **(B)** daily mean temperatures, **(C)** daily maximum temperatures, and **(D)** daily minimum temperatures. P-values indicate significant differences in temperature metrics between experimental treatments from an ANOVA, and eta-squared ($\eta^2$) indicates the effect size of the treatment term in the ANOVA.

**Table S1. *In situ* site metadata and *ex situ* experimental treatment conditions.** (**A**) Site ID, type, and GPS coordinates for sites where *Siderastrea siderea* (N=9 colonies/site) were collected across the Bocas del Toro Reef Complex, Panamá (SP=STRI Point, PD=Punta Donato, CI=Cristobal Island, CA=Cayo de Agua, BN=Bastimentos North, and BS=Bastimentos South). Average, minimum, and maximum daily temperature parameters (˚C) are reported for four sites where temperature loggers were recovered (SP, PD, CI, and CA). When error is shown, values represent means ± standard error (SE). When no error is shown, values represent maximum values. (**B**) All values are reported as overall means of the daily mean ± SE of the temperature parameters throughout the 50-day diel temperature variability (DTV) treatments.

| **A) *In situ* site metadata** | | | | | | | | |
|---|---|---|---|---|---|---|---|---|
| **Site** | **Type** | **GPS** | **Daily Mean** | **Daily Min** | **Daily Max** | **DTV** | **Max DTV** | **Overall Max** |
| SP | Inshore | 9.352434, -82.264553 | 29.32 ± 0.04°C | 29.08 ± 0.04°C | 29.62 ± 0.04°C | 0.54 ± 0.01°C | 1.26°C | 31.59°C |
| PD | Inshore | 9.36924, -82.364037 | 29.70 ± 0.04°C | 29.36 ± 0.04°C | 30.07 ± 0.04°C | 0.71 ± 0.01°C | 1.40°C | 32.15°C |
| CI | Inshore | 9.2654486, -82.243511 | 29.86 ± 0.04°C | 29.37 ± 0.04°C | 30.54 ± 0.05°C | 1.17 ± 0.03°C | 3.17°C | 33.55°C |
| CA | Offshore | 9.193858, -82.053951 | 29.45 ± 0.04°C | 28.99 ± 0.04°C | 29.97 ± 0.05°C | 0.98 ± 0.02°C | 2.41°C | 32.43°C |
| BN | Offshore | 9.348495, -82.176505 | N/A | N/A | N/A | N/A | N/A | N/A |
| BS | Offshore | 9.287438, -82.092315 | N/A | N/A | N/A | N/A | N/A | N/A |

| **B) *Ex situ* experimental conditions (Fig. S10)** | | | | |
|---|---|---|---|---|
| **Treatment** | **Variability** | **Mean** | **Maximum** | **Minimum** |
| Control | 0.43 ± 0.04 | 29.45°C ± 0.01 | 29.59°C ± 0.02 | 29.16°C ± 0.04 |
| DTV | 2.88°C ± 0.02 | 29.40°C ± 0.02 | 31.28°C ± 0.03 | 28.40°C ± 0.02 |

**Table S2.** Betadispersion and PERMANOVA results from bacterial community data (all ASVs), presented in Figs. 1B and S6A,B. Explicit models are listed under each section for the data being considered. Sum Sq=sum of squares, DF=degrees of freedom, $R^2$=percentage of variance explained.

| **Bacterial Community Spatial Statistics (Figs. 1B, S6)** | | | | | |
|---|---|---|---|---|---|
| ***Baseline Dispersion*** | | | | | |
| **Factor/Comparison** | **DF** | **Sum Sq** | **Mean Sq** | **F-value** | **P-value** |
| | **Model = betadisper(distance.taxa ~ lineage)** | | | | |
| Lineage | 1 | 0.000073 | 0.000073 | 0.036 | 0.85 |
| ***Baseline Spatial Structure*** | | | | | |
| **Factor/Comparison** | **DF** | **Sum Sq** | **$R^2$** | **F-Value** | **P-value** |
| | **Model = adonis(taxa ~ lineage, permutations = 999)** | | | | |
| Lineage | 1 | 0.60 | 0.031 | 1.46 | **0.003** |
| ***Post-DTV Dispersion*** | | | | | |
| **Factor/Comparison** | **DF** | **Sum Sq** | **Mean Sq** | **F-value** | **P-value** |
| | **Model = betadisper(distance.taxa ~ lineage)** | | | | |
| Lineage | 1 | 0.0062 | 0.0062 | 4.57 | **0.036** |
| | **Model = betadisper(distance.taxa ~ DTV treatment)** | | | | |
| DTV | 1 | 0.00055 | 0.00055 | 0.35 | 0.55 |
| ***Post-DTV Spatial Structure*** | | | | | |
| **Factor/Comparison** | **DF** | **Sum Sq** | **$R^2$** | **F-Value** | **P-value** |
| | **Model = adonis(taxa ~ DTV treatment + lineage, permutations = 999)** | | | | |
| DTV Treatment | 1 | 0.61 | 0.02 | 1.4 | **0.045** |
| Lineage | 1 | 0.54 | 0.018 | 1.25 | 0.11 |

**Table S3.** ANOVA results from bacterial community diversity data, including separate analyses for Shannon Index, Simpson Index, OTU Richness, and Evenness for baseline and post-DTV timepoints, presented in Figs. S2 and S6. Explicit models are listed under each section for the data being considered. Sum Sq=sum of squares, DF=degrees of freedom.

| **Bacterial Community Diversity Statistics (Figs. S2 and S6)** | | | | | |
|---|---|---|---|---|---|
| **Shannon Index** | | | | | |
| **Factor/Comparison** | **DF** | **Sum Sq** | **Mean Sq** | **F-value** | **P-value** |
| ***Baseline*** | **Model = aov(Shannon ~ lineage)** | | | | |
| Lineage | 1 | 1.55 | 1.55 | 2.37 | 0.13 |
| ***Post-DTV*** | **Model = aov(Shannon ~ DTV treatment + lineage)** | | | | |
| Lineage | 1 | 1.05 | 1.05 | 0.96 | 0.33 |
| DTV Treatment | 1 | 1.01 | 1.006 | 0.92 | 0.34 |
| **Simpson Index** | | | | | |
| ***Baseline*** | **Model = aov(Simpson ~ lineage)** | | | | |
| Lineage | 1 | 5406 | 5406 | 3.05 | 0.09 |
| ***Post-DTV*** | **Model = aov(Simpson ~ DTV treatment + lineage)** | | | | |
| Lineage | 1 | 93 | 93.31 | 0.316 | 0.576 |
| DTV Treatment | 1 | 108 | 108.1 | 0.366 | 0.547 |
| **OTU Richness** | | | | | |
| ***Baseline*** | **Model = aov(Richness ~ lineage)** | | | | |
| Lineage | 1 | 8431 | 8431 | 0.355 | 0.55 |
| ***Post-DTV*** | **Model = aov(Richness ~ DTV treatment + lineage)** | | | | |
| DTV Treatment | 1 | 66991 | 66991 | 5.4 | **0.02** |
| Lineage | 1 | 9 | 9 | 0.001 | 0.98 |
| **Evenness** | | | | | |
| ***Baseline*** | **Model = aov(Evenness ~ lineage)** | | | | |
| Lineage | 1 | 0.029 | 0.029 | 3.72 | 0.06 |
| ***Post-DTV*** | **Model = aov(Evenness ~ DTV treatment + lineage)** | | | | |
| Lineage | 1 | 0.029 | 0.029 | 1.01 | 0.32 |
| DTV Treatment | 1 | 0.044 | 0.044 | 1.56 | 0.22 |

**Table S4.** PERMANOVA results from holobiont phenome data presented in Fig. 2A and Fig. 3D. Explicit models are listed under each section for the time point being considered (baseline and end of experiment). Sum Sq=sum of squares, Mean Sq=mean square of the error, DF=degrees of freedom, $R^2$=percentage of variance explained, Omega-sq=effect size. N=42 for baseline and N=46 for end of experiment.

| **Holobiont Phenome Physiology PERMANOVA Statistics** | | | | | | | |
|---|---|---|---|---|---|---|---|
| **Factor/ Comparison** | **DF** | **Sum Sq** | **Mean Sq** | **F-Value** | **$R^2$** | **Omega-sq** | **P-value** |
| ***Baseline*** | **Model = adonis(scores ~ lineage)** | | | | | | |
| Lineage | 1 | 0.008 | 0.008 | 7.65 | 0.16 | 0.137 | **3e-04** |
| ***End of Experiment*** | **Model = adonis(scores ~ lineage + dtv_treatment)** | | | | | | |
| Lineage | 1 | 0.036 | 0.036 | 12.57 | 0.18 | 0.201 | **2e-04** |
| DTV Treatment | 1 | 0.034 | 0.034 | 11.94 | 0.18 | 0.19 | **3e-04** |

**Table S5.** Linear model and linear mixed model results for individual host and symbiont phenotype data presented in Fig. S4. Each physiology metric includes separate models for the two time points considered: 1) before acclimation (Baseline) and 2) at the end of experiment (End of Experiment). Explicit models are listed under each section for the physiology metric and time point being considered. Sum Sq=sum of squares, Mean Sq=mean square of the error, DF=degrees of freedom.

| **Individual Physiology Metric Statistics (Fig. S4)** | | | | | |
|---|---|---|---|---|---|
| **Factor** | **DF** | **Sum Sq** | **Mean Sq** | **F-value** | **P-value** |
| ***Total Protein, Baseline*** | | | | | **Model = lm(protein ~ lineage)** |
| Lineage | 1 | 0.079 | 0.079 | 12.00 | **0.001** |
| ***Total Protein, End of Experiment*** | | | | | **Model = lmer(protein ~ lineage + dtv_treatment + 1\|genet)** |
| DTV treatment | 1 | 0.0036 | 0.0036 | 0.39 | 0.54 |
| Lineage | 1 | 0.11 | 0.11 | 11.84 | **0.002** |
| ***Host Carbohydrate, Baseline*** | | | | | **Model = lm(host_carb ~ lineage)** |
| Lineage | 1 | 0.052 | 0.052 | 7.14 | **0.01** |
| ***Host Carbohydrate, End of Experiment*** | | | | | **Model = lmer(host_carb ~ lineage + dtv_treatment + 1\|genet)** |
| Lineage | 1 | 0.298 | 0.298 | 16.46 | **0.0002** |
| DTV treatment | 1 | 0.372 | 0.372 | 20.55 | **5.4e-05** |
| ***Symbiont Carbohydrate, Baseline*** | | | | | **Model = lm(sym_carb ~ lineage)** |
| Lineage | 1 | 0.022 | 0.022 | 9.01 | **0.005** |
| ***Symbiont Carbohydrate, End of Experiment*** | | | | | **Model = lmer(sym_carb ~ lineage + dtv_treatment + 1\|genet)** |
| Lineage | 1 | 0.012 | 0.012 | 3.20 | 0.08 |
| DTV treatment | 1 | 0.049 | 0.049 | 12.53 | **0.0009** |
| ***Chlorophyll a, Baseline*** | | | | | **Model = lm(chla ~ lineage)** |
| Lineage | 1 | 297.5 | 297.5 | 20.45 | **5.14e-05** |
| ***Chlorophyll a, End of Experiment*** | | | | | **Model = lmer(sym_carb ~ lineage + dtv_treatment + 1\|genet)** |

| | | | | | |
|---|---|---|---|---|---|
| DTV treatment | 1 | 0.005 | 0.005 | 0.0016 | 0.97 |
| Lineage | 1 | 29.06 | 29.06 | 9.72 | **0.003** |
| ***Symbiont Density, Baseline*** | **Model = lm(syms ~ lineage)** | | | | |
| Lineage | 1 | 9.37e12 | 9.37e12 | 18.37 | **0.0002** |
| ***Symbiont Density, End of Experiment*** | **Model = lmer(syms ~ lineage + dtv_treatment + 1\|genet)** | | | | |
| Lineage | 1 | 3.08e12 | 3.08e12 | 11.74 | **0.001** |
| DTV treatment | 3 | 1.76e12 | 1.76e12 | 6.72 | **0.0096** |

**Table S6.** PERMANOVA results from skeleton morphology data presented in Fig. 2B. Explicit model is listed. Sum Sq=sum of squares, Mean Sq=mean square of the error, DF=degrees of freedom, $R^2$=percentage of variance explained, Omega-sq=effect size. N=42.

| **Skeleton Morphology PERMANOVA Statistics (Fig. 2B)** | | | | | | | |
|---|---|---|---|---|---|---|---|
| **Factor** | **DF** | **Sum Sq** | **Mean Sq** | **F-Value** | **$R^2$** | **Omega-sq** | **P-value** |
| | **Model = adonis(scores ~ lineage + site_of_origin)** | | | | | | |
| Lineage | 1 | 0.017 | 0.017 | 10.08 | 0.20 | 0.177 | **0.0007** |

**Table S7.** Linear model results for corallite area and light enhancement factor (LEF) differences across lineages, presented in Fig. 2D and Fig. S5. The explicit model is listed. DF=degrees of freedom, SE=standard error. N=45 for corallite area from all sites (L1 N=29, L2 N=16), N=8 for corallite area from Cristobal Island [CI] only (L1 N=4, L2 N=4). N=42 for LEF from all sites (L1 N=27, L2 N=15), N=7 for LEF from CI only (L1 N=4, L2 N=3).

| **Corallite Area – All Sites (Fig. S5A)** | | | | |
|---|---|---|---|---|
| **Factor/Comparison** | **DF** | **SE** | **T-value** | **P-value** |
| | | | | **Model = lm(corallite_area ~ lineage)** |
| Intercept | 8.8205 | 0.3375 | 26.134 | **<2e-16** |
| Lineage (L2) | 5.0749 | 0.566 | 8.966 | **2.13e-11** |
| **Corallite Area – Cristobal Island Only (Fig. S5B)** | | | | |
| **Factor/Comparison** | **Estimate** | **SE** | **T-value** | **P-value** |
| | | | | **Model = lm(corallite_area ~ lineage)** |
| Intercept | 7.52 | 0.73 | 10.4 | **4.73e-05** |
| Lineage (L2) | 6.01 | 1.03 | 5.9 | **0.001** |
| **Light Enhancement Factor (LEF) – All Sites (Figure 2D)** | | | | |
| **Factor/Comparison** | **Estimate** | **SE** | **T-value** | **P-value** |
| | | | | **Model = lm(lef ~ lineage)** |
| Intercept | 1.34 | 0.062 | 21.42 | **<2e-16** |
| Lineage (L2) | 0.80 | 0.104 | 7.7 | **2.01e-09** |
| **Light Enhancement Factor (LEF) – Cristobal Island Only (Figure S5C)** | | | | |
| **Factor/Comparison** | **Estimate** | **SE** | **T-value** | **P-value** |
| | | | | **Model = lm(lef ~ lineage)** |
| Intercept | 1.27 | 0.16 | 7.89 | **0.0005** |
| Lineage (L2) | 0.68 | 0.246 | 2.76 | **0.04** |

**Table S8.** Linear model (summarized using the *anova*() function) and Tukey's HSD results for coral growth data presented in Figs. 3A,B. Explicit models are listed under each section for the time point being considered (during DTV, or during thermal stress and recovery). Sample sizes are as follows during variability: Control (L1=24, L2=15), DTV (L1=25, L2=18). Sample sizes are as follows during thermal stress and recovery: Control (L1=24, L2=13), DTV (L1=25, L2=17).

| **Coral Growth Statistics (Fig. 3A,B)** | | | | | |
|---|---|---|---|---|---|
| **Factor/Comparison** | **Sum Sq** | **Mean Sq** | **DF** | **F-value** | **P-value** |
| ***During DTV*** | **Model = lmer(growth ~ treatment + lineage + (1\|genet))** | | | | |
| Treatment | 1.46e-07 | 1.46e-07 | 1 | 5.66 | **0.022** |
| Lineage | 8.47e-08 | 8.47e-08 | 1 | 3.28 | 0.077 |
| ***Tukey HSD*** | **Estimate** | **SE** | **DF** | **T-value** | **P-value** |
| Control (L1-L2) | 8.42e-05 | 7.32e-05 | 78.7 | 1.15 | 0.25 |
| DTV (L1-L2) | 1.30e-04 | 6.95e-05 | 74.3 | 1.87 | 0.13 |
| ***During Thermal Stress + Recovery*** | **Model = lmer(growth ~ treatment + lineage + (1\|genet))** | | | | |
| Treatment | 1.11e-07 | 1.11e-07 | 1 | 1.25 | 0.27 |
| Lineage | 1.13e-06 | 1.13e-06 | 1 | 12.77 | **0.0008** |
| ***Tukey HSD*** | **Estimate** | **SE** | **DF** | **T-value** | **P-value** |
| Control (L1-L2) | 0.00033 | 0.00013 | 80.9 | 2.56 | **0.012** |
| DTV (L1-L2) | 0.00036 | 0.00012 | 77.3 | 3.13 | **0.005** |

**Table S9.** Linear model (summarized using the *anova*() function) and Tukey HSD (within timepoints across lineages) results for photochemical efficiency (Fv/Fm) data presented in Fig. 3C. Explicit model is listed. Sample sizes for each treatment are as follows: Control (N=27), DTV (N=29).

| **Photochemical Efficiency (Fv/Fm) Statistics (Fig. 3C)** | | | | | |
|---|---|---|---|---|---|
| **Factor/Comparison** | **Sum Sq** | **Mean Sq** | **DF** | **F-value** | **P-value** |
| | **Model = lmer(fvfm ~ time*lineage*dtv_treatment + 1\|genet)** | | | | |
| Time | 0.035 | 0.006 | 6 | 4.11 | **0.0006** |
| Lineage | 0.01 | 0.01 | 1 | 6.82 | **0.013** |
| DTV Treatment | 0.002 | 0.002 | 1 | 1.61 | 0.206 |
| Time*Lineage | 0.014 | 0.002 | 6 | 1.66 | 0.13 |
| Time*DTV Treatment | 0.006 | 0.0009 | 6 | 0.67 | 0.67 |
| Lineage*DTV Treatment | 0.002 | 0.002 | 1 | 1.60 | 0.21 |
| Time*Lineage*DTV Treatment | 0.008 | 0.001 | 6 | 0.99 | 0.43 |
| ***Tukey HSD*** | **Estimate** | **SE** | **DF** | **T-value** | **P-value** |
| Time 1 (L1-L2) | 0.025 | 0.0197 | 72.9 | 1.29 | 0.202 |
| Time 2 (L1-L2) | 0.045 | 0.0197 | 72.9 | 2.29 | 0.058 |
| Time 3 (L1-L2) | 0.061 | 0.0197 | 72.9 | 3.098 | **0.011** |
| Time 4 (L1-L2) | 0.060 | 0.0197 | 72.9 | 3.063 | **0.011** |
| Time 5 (L1-L2) | 0.04 | 0.0197 | 72.9 | 2.043 | 0.078 |
| Time 6 (L1-L2) | 0.034 | 0.0197 | 72.9 | 1.74 | 0.10 |
| Time 7 (L1-L2) | 0.036 | 0.0197 | 72.9 | 1.84 | 0.099 |

**Table S10.** Linear model results for historical growth differences across lineages. Two time intervals are included: 1) recent data (1980-2014; Fig. 4) and 2) all data (1880-2014; Fig. S8). The explicit model is listed. N=24 for both time intervals (L1 N=16, L2 N=8).

| | | | | |
|---|---|---|---|---|
| ***Linear Extension (1980-2014)*** | | | | |
| **Factor/Comparison** | **Estimate** | **SE** | **T-value** | **P-value** |
| **Model = lm(linear_extension ~ lineage)** | | | | |
| Intercept | 0.516 | 0.033 | 15.84 | **1.64e-13** |
| Lineage (L2) | -0.13 | 0.056 | -2.38 | **0.026** |
| ***Linear Extension (All Years)*** | | | | |
| **Factor/Comparison** | **Estimate** | **SE** | **T-value** | **P-value** |
| **Model = lm(linear_extension ~ lineage)** | | | | |
| Intercept | 0.52 | 0.032 | 16.25 | **9.68e-14** |
| Lineage (L2) | -0.13 | 0.056 | -2.37 | **0.027** |
| ***Skeletal Density (1980-2014)*** | | | | |
| **Factor/Comparison** | **Estimate** | **SE** | **T-value** | **P-value** |
| **Model = lm(density ~ lineage)** | | | | |
| Intercept | 1.30 | 0.03 | 44.07 | **<2e-16** |
| Lineage (L2) | 0.16 | 0.05 | 3.19 | **0.004** |
| ***Skeletal Density (All Years)*** | | | | |
| **Factor/Comparison** | **Estimate** | **SE** | **T-value** | **P-value** |
| **Model = lm(density ~ lineage)** | | | | |
| Intercept | 1.31 | 0.03 | 43.09 | **<2e-16** |
| Lineage (L2) | 0.16 | 0.053 | 2.96 | **0.007** |

| ***Calcification (1980-2014)*** | | | | |
|---|---|---|---|---|
| **Factor/Comparison** | **Estimate** | **SE** | **T-value** | **P-value** |
| | | | | **Model = lm(calcification ~ lineage)** |
| Intercept | 0.67 | 0.04 | 16.59 | **6.38e-14** |
| Lineage (L2) | -0.12 | 0.07 | -1.68 | 0.11 |
| ***Calcification (All Years)*** | | | | |
| **Factor/Comparison** | **Estimate** | **SE** | **T-value** | **P-value** |
| | | | | **Model = lm(corallite_area ~ lineage)** |
| Intercept | 0.68 | 0.04 | 17.12 | **3.34e-14** |
| Lineage (L2) | -0.12 | 0.069 | -1.72 | 0.1 |

**Table S11.** Linear model results for the relationship between time and historical temperature data from the HadISST dataset (both summer mean and annual mean temperature), including data from 1870-2023.

| ***Summer Mean Temperature*** | | | | |
|---|---|---|---|---|
| **Factor/Comparison** | **Estimate** | **SE** | **T-value** | **P-value** |
| **Model = lm(Summer_Mean ~ Year)** | | | | |
| Intercept | 67.32 | 161.2 | 0.418 | 0.68 |
| Summer Mean | 66.78 | 5.73 | 11.66 | **<2e-16** |
| ***Annual Mean Temperature*** | | | | |
| **Factor/Comparison** | **Estimate** | **SE** | **T-value** | **P-value** |
| **Model = lm(Annual_Mean ~ Year)** | | | | |
| Intercept | -272.7 | 172.0 | -1.59 | 0.12 |
| Annual Mean | 79.31 | 6.15 | 12.9 | **<2e-16** |

**Table S12.** Linear model results for the relationship between historical calcification derived from coral cores and historical temperature data from the HadISST dataset (both summer mean and annual mean temperature) across lineages. Two time intervals are included: 1) recent data (1980-2014) and 2) all data (1880-2014) (Fig. S9C-F). The explicit model is listed. N=24 for both time intervals (L1 N=16, L2 N=8).

| ***Summer Mean Temperature (1980-2014)*** | | | | |
|---|---|---|---|---|
| **Factor/Comparison** | **Estimate** | **SE** | **T-value** | **P-value** |
| | | | | **Model = lm(Calcification ~ summer_mean + lineage)** |
| Summer Mean | 0.0034 | 0.030 | 0.113 | 0.910 |
| Lineage (L2) | -0.13 | 0.018 | -7.34 | **5.29e-13** |
| ***Summer Mean Temperature (All Years)*** | | | | |
| **Factor/Comparison** | **Estimate** | **SE** | **T-value** | **P-value** |
| | | | | **Model = lm(Calcification ~ summer_mean + lineage)** |
| Summer Mean | 0.02 | 0.018 | 1.16 | 0.25 |
| Lineage (L2) | -0.11 | 0.014 | -7.61 | **5.44e-14** |
| ***Annual Mean Temperature (1980-2014)*** | | | | |
| **Factor/Comparison** | **Estimate** | **SE** | **T-value** | **P-value** |
| | | | | **Model = lm(Calcification ~ annual_mean + lineage)** |
| Annual Mean | -0.018 | 0.035 | -0.52 | 0.60 |
| Lineage (L2) | -0.13 | 0.018 | -7.34 | **5.51e-13** |
| ***Annual Mean Temperature (All Years)*** | | | | |
| **Factor/Comparison** | **Estimate** | **SE** | **T-value** | **P-value** |
| | | | | **Model = lm(Calcification ~ annual_mean + lineage)** |
| Annual Mean | 0.018 | 0.021 | 0.084 | 0.40 |

| | | | | |
|---|---|---|---|---|
| Lineage (L2) | -0.11 | 0.014 | -7.59 | **6.21e-14** |

**Table S13. ANOVA and Tukey's HSD results for daily *in situ* temperature data.** Explicit models are listed under each section for the metric being considered. Sum Sq=sum of squares, Mean Sq=mean square of the error, DF=degrees of freedom. For Tukey HSD outputs, estimate, lower, and upper columns represent the difference between the compared mean values and 95% confidence intervals. Punta Donato=PD, STRI Point=SP, Cristobal Island=CI, Bastimentos North=BN, Bastimentos South=BS, Cayo de Agua=CA.

| **Daily *in situ* Temperature Statistics (Fig. S2B)** | | | | | |
|---|---|---|---|---|---|
| **Factor/Comparison** | **DF** | **Sum Sq** | **Mean Sq** | **F-value** | **P-value** |
| ***Daily Range*** | **Model = aov(DailyRange ~ site of origin)** | | | | |
| | 3 | 100.2 | 33.42 | 239.1 | **<2e-16** |
| ***Tukey HSD*** | | **Estimate** | **Lower** | **Upper** | **P-value** |
| SP-PD | | -0.167 | -0.232 | -0.101 | **<0.0001** |
| CI-PD | | 0.459 | 0.394 | 0.524 | **<0.0001** |
| CA-PD | | 0.270 | 0.205 | 0.336 | **<0.0001** |
| CI-SP | | 0.626 | 0.561 | 0.692 | **<0.0001** |
| CA-SP | | 0.437 | 0.372 | 0.503 | **<0.0001** |
| CA-CI | | -0.189 | -0.255 | -0.124 | **<0.0001** |
| ***Daily Mean*** | **Model = aov(DailyMean ~ site of origin)** | | | | |
| | 3 | 77.9 | 25.98 | 39.26 | **<0.0001** |
| ***Tukey HSD*** | | **Estimate** | **Lower** | **Upper** | **P-value** |
| SP-PD | | -0.388 | -0.530 | -0.245 | **<0.0001** |
| CI-PD | | 0.159 | 0.016 | 0.301 | **0.022** |
| CA-PD | | -0.250 | -0.393 | -0.108 | **<0.0001** |
| CI-SP | | 0.547 | 0.404 | 0.689 | **<0.0001** |
| CA-SP | | 0.138 | -0.005 | 0.280 | 0.063 |
| CA-CI | | -0.409 | -0.552 | -0.267 | **<0.0001** |

**Table S14.** ANOVA and Tukey's HSD results for *ex situ* experiment temperature data presented in Fig. S10. Explicit models are listed under each section for the temperature metric being considered. Sum Sq=sum of squares, Mean Sq=mean square of the error, DF=degrees of freedom, $\eta^2$=Eta-squared.

| **Daily DTV Treatment Temperature Statistics (Fig. S10)** | | | | | | |
|---|---|---|---|---|---|---|
| **Factor/Comparison** | **DF** | **Sum Sq** | **Mean Sq** | **F-value** | $\eta^2$ | **P-value** |
| ***Daily Range*** | **Model = aov(DailyRange ~ treatment)** | | | | | |
| DTV Treatment | 1 | 150.4 | 150.4 | 2823 | 0.97 | **<2e-16** |
| **Factor/Comparison** | **DF** | **Sum Sq** | **Mean Sq** | **F-value** | $\eta^2$ | **P-value** |
| ***Daily Mean*** | **Model = aov(DailyMean ~ treatment)** | | | | | |
| DTV Treatment | 1 | 0.052 | 0.052 | 4.28 | 0.04 | **0.04** |
| **Factor/Comparison** | **DF** | **Sum Sq** | **Mean Sq** | **F-value** | $\eta^2$ | **P-value** |
| ***Daily Minimum*** | **Model = aov(DailyMin ~ treatment)** | | | | | |
| DTV Treatment | 1 | 14.52 | 14.52 | 266.3 | 0.73 | **<2e-16** |
| **Factor/Comparison** | **DF** | **Sum Sq** | **Mean Sq** | **F-value** | $\eta^2$ | **P-value** |
| ***Daily Maximum*** | **Model = aov(DailyMax ~ treatment)** | | | | | |
| DTV Treatment | 1 | 71.47 | 71.47 | 3021 | 0.97 | **<2e-16** |
